# De novo production of an antitumor precursor actinocin and other medicinal molecules from kynurenine pathway in *Escherichia coli*

**DOI:** 10.1101/2023.07.31.551311

**Authors:** Komal Sharma, Mohammad Rifqi Ghiffary, GaRyoung Lee, Hyun Uk Kim

## Abstract

Kynurenine pathway has a potential to convert L-tryptophan into multiple medicinal molecules. This study aims to explore the biosynthetic potential of kynurenine pathway for the efficient production of actinocin, an antitumor precursor selected as a proof-of-concept target molecule. Kynurenine pathway is first constructed in *Escherichia coli* by testing various combinations of biosynthetic genes from four different organisms. Metabolic engineering strategies are next performed to improve the production by inhibiting a competing pathway, and enhancing intracellular supply of a cofactor *S*-adenosyl-L-methionine, and ultimately to produce actinocin from glucose. Metabolome analysis further suggests additional gene overexpression targets, which finally leads to the actinocin titer of 719 mg/L. *E. coli* strain engineered to produce actinocin is further successfully utilized to produce 350 mg/L of kynurenic acid, a neuroprotectant, and 1401 mg/L of 3-hydroxyanthranilic acid, an antioxidant, also from glucose. These competitive production titers demonstrate the biosynthetic potential of kynurenine pathway as a source of multiple medicinal molecules. The approach undertaken in this study can be useful for producing other molecules associated with kynurenine pathway.

## Introduction

Kynurenine pathway is responsible for converting L-tryptophan into several biologically active molecules besides nicotinamide adenine dinucleotide (NAD^+^)^1^. Here, kynurenine pathway deserves attention for producing multiple medicinal molecules because its intermediates may serve as a drug or a direct drug precursor (Fig. 1 and Supplementary Fig. 1). For example, kynurenic acid is a neuroprotective and anticonvulsant metabolite^2^, and has also been suggested as a dietary supplementation^3^ and a weight-loss agent^4^. 3-Hydroxyanthranilic acid (3-HA) is another metabolite from kynurenine pathway with antioxidant^5^ and anti-aging properties^6^, and plays anti-inflammatory and neuroprotective roles during inflammation^7^. While the kynurenine pathway has been extensively studied in mammals, it is also identified in several microorganisms, including *Saccharomyces cerevisiae*^8^ and *Streptomyces* species (Supplementary Fig. 1). Examples of *Streptomyces* species include *Streptomyces coelicolor*, *Streptomyces parvulus*, *Streptomyces anulatus* and *Streptomyces avermitilis* where kynurenine pathway plays a role in the biosynthesis of secondary metabolites, such as antibiotics (e.g., daptomycin^9–11^, taromycin^12^ and triostin^13^) and antitumor agents (e.g., echinomycin^14^, thiocoraline^15^ and actinomycins^16^). Thus, the kynurenine pathway has a potential for generating multiple medicinal molecules, and also at their high-production level, but it has not been extensively studied for its biosynthetic potential.

**Fig. 1.**
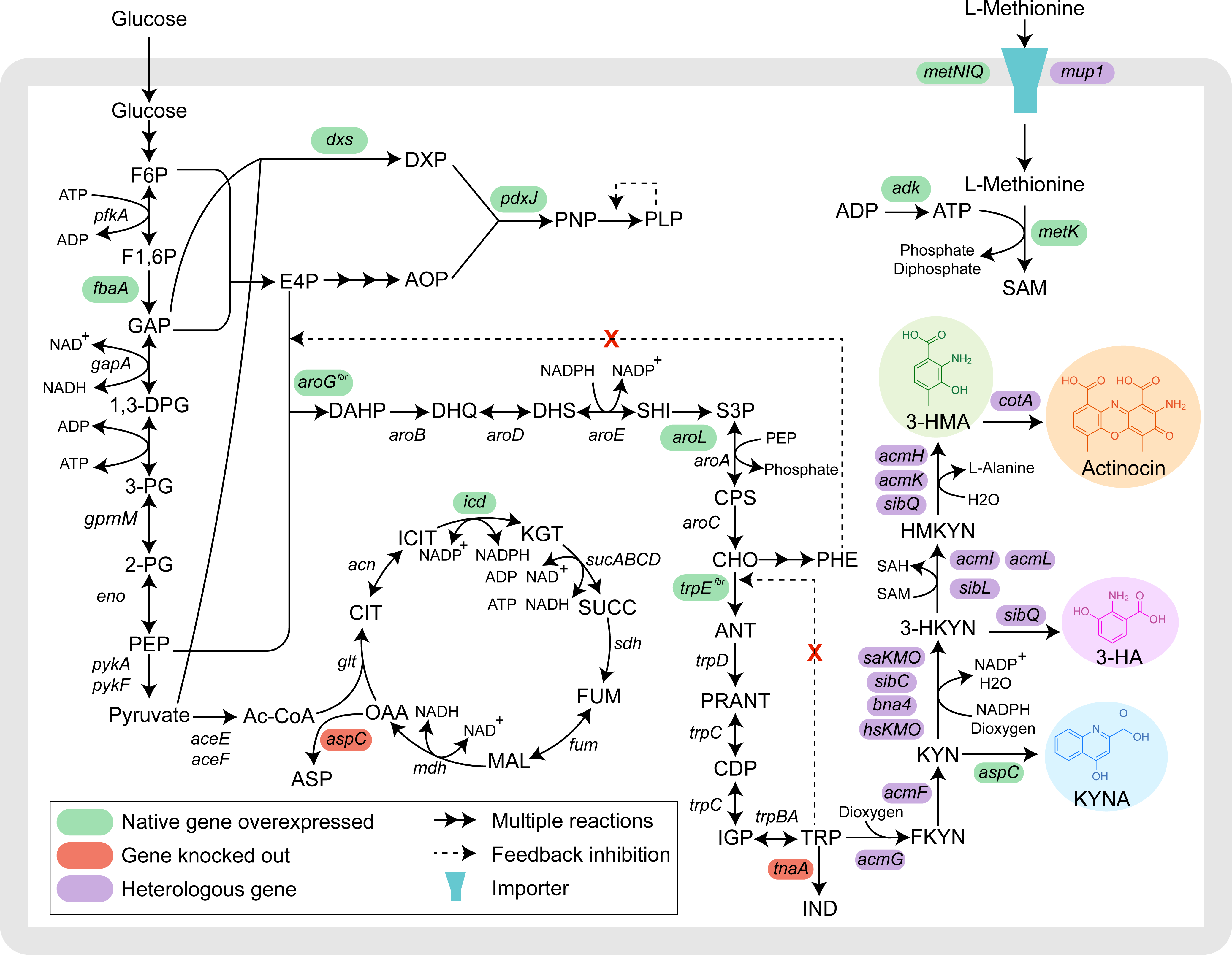
Overall metabolic engineering strategies for the de novo production of three medicinal molecules from kynurenine pathway. For the construction of kynurenine pathway, following genes were examined in this study: tryptophan 2,3-dioxygenase (*acmG*), kynurenine formamidase (*acmF*), kynurenine 3-monooxygenase (*saKMO*, *sibC*, *bna4* and *hsKMO*), methyltransferase (*acmI*, *acmL* and *sibL*), and kynureninase (*acmH*, *acmK*, and *sibQ*). Laccase (*cotA*) was additionally expressed to produce actinocin. For the enhanced production of 3-hydroxy-4-methylanthranilic acid (3-HMA), a precursor of actinocin, following were examined: knockout of *aspC* encoding aspartate aminotransferase to avoid the competition with kynurenine pathway; overexpression of genes encoding native methionine transporter genes (*metNIQ*), heterologous methionine permease (*mup1*), native *S*-adenosylmethionine synthase (*metK*), and/or adenylate kinase (*adk*) to enhance the biosynthesis of *S*-adenosyl-L-methionine (SAM), a cofactor required for 3-HMA; overexpression of genes encoding 1-deoxy-D-xylulose-5-phosphate synthase (*dxs*) and pyridoxine 5’-phosphate synthase (*pdxJ*) to enhance the biosynthesis of pyridoxal 5’-phosphate (PLP), the other cofactor required for 3-HMA. For the enhanced production of actinocin, following were additionally conducted: overexpression of genes that encode feedback-resistant 3-deoxy-D-arabino-heptulosonate 7-phosphate synthase (*aroG^fbr^*), shikimate kinase II (*aroL*), and feedback-resistant anthranilate synthase (*trpE^fbr^*); knockout of *tnaA* encoding tryptophanase; and overexpression of genes that encode fructose 1,6-bisphosphate aldolase (*fbaA*) and isocitrate dehydrogenase (*icd*). Abbreviations of the metabolites are: 1,3-DPG 1,3-diphospho-D-glycerate; 2-PG, 2-phosphoglycerate; 3-HA, 3-hydroxyanthranilic acid; 3-HKYN, 3-hydroxykynurenine; 3-HMA, 3-hydroxy-4-methyl anthranilic acid; 3-PG, 3-phosphoglycerate; Ac-CoA, acetyl-Coenzyme A; ANT, anthranilate; ASP, aspartate; AOP, 3-amino-2-oxopropyl phosphate; CDP, 1-(2-carboxyphenylamino)-1-deoxy-D-ribulose-5-phosphate; CHO, chorismate; CIT, citrate; CPS, 5-*o*-(1-carboxyvinyl)-3-phosphoshikimate; DAHP, 3-deoxy-arabino-heptulonate-7-phosphate; DHQ, 3-dehydroquinate; DHS, 3-dehydroshikimate; DXP, deoxyxylulose 5-phosphate; E4P, erythrose 4-phosphate; F1,6P, fructose 1,6-bisphosphate; F6P, fructose-6-phosphate; FKYN, *N*-formyl-L-kynurenine; FUM, fumarate; GAP, glyceraldehyde 3-phosphate; HMKYN, 3-hydroxy-4-methylkynurenine; ICIT, isocitrate; IGP, indole 3-glycerol phosphate; IND, indole; KGT, α-ketoglutarate; KYN, L-kynurenine; MAL, malate; OAA, oxaloacetate; PEP, phosphoenolpyruvate; PLP, pyridoxal 5’-phosphate; PNP, pyridoxine 5’-phosphate; PRANT, *N*-(5-phospho-D-ribosyl) anthranilate; S3P, shikimate 3-phosphate; SAH, *S*-adenosyl-L-homocysteine; SAM, *S*-adenosyl-L-methionine; SUCC, succinate; SHI, shikimate; TRP, tryptophan.

In this study, we optimally express kynurenine pathway in *Escherichia coli*, and examine the biosynthetic potential of this pathway by producing three medicinal molecules as proof-of-concept targets. Our primary objective molecule is actinocin, which is the phenoxazinone chromophore of an antitumor agent actinomycin D^17, 18^. We additionally attempted to produce kynurenic acid and 3-HA. For the production of actinocin, we first constructed kynurenine pathway in *E. coli* for the enhanced production of 3-hydroxy-4-methylanthranilic acid (3-HMA), a precursor of actinocin. 3-HMA is also a precursor of several antitumor agents, including sibiromycin^19, 20^ and anthramycin^21^. Heterologous expression of the kynurenine pathway was optimized by examining multiple combinations of biosynthetic genes from *S. anulatus*, *Streptosporangium sibiricum*, *S. cerevisiae*, and *Homo sapiens*. To further improve the 3-HMA biosynthesis, metabolic engineering was performed to inhibit a competing pathway, and to enhance the intracellular supply of *S*-adenosyl-L-methionine (SAM) and pyridoxal 5’-phosphate (PLP), both required as cofactors for the biosynthesis of 3-HMA. Best performing strain BAP1Δ*aspC* with pHMA9, pSAM3 and pACN1 was further engineered to produce actinocin from glucose de novo. Subsequent metabolome analysis suggested two nonintuitive gene manipulation targets, which further improved the actinocin titer by more than twofold (i.e., 354 mg/L to 719 mg/L) via fed-batch fermentation. BAP1Δ*aspC*Δ*tnaA* strain initially constructed for actinocin was also transformed with different plasmids to produce two additional molecules, kynurenic acid and 3-HA, both associated with kynurenine pathway (Fig. 1).

To our knowledge, this study is the first to intensively examine the biosynthetic potential of kynurenine pathway for producing multiple medicinal molecules at competitive level. Based on the resulting competitive production titers of actinocin, kynurenic acid and 3-HA, this study demonstrates the biosynthetic value of kynurenine pathway. In particular, the approach undertaken in this study can be useful for efficiently producing other kynurenine pathway-derived molecules that may have a wide range of applications, especially drugs (e.g., questiomycin^22^) and also dyes (e.g., ommatin D^23^ and cinnabarins^24, 25^).

## Results

### Heterologous expression of kynurenine pathway for the biosynthesis of 3-HMA

Kynurenine pathway was first constructed in *E. coli* by expressing heterologous biosynthetic genes for 3-HMA (Fig. 1 and Supplementary Tables 1-4). Biosynthesis of 3-HMA requires five enzymatic steps from L-tryptophan. Initial target was to biosynthesize L-kynurenine by expressing two heterologous genes *acmG* and *acmF*, which encode tryptophan 2,3-dioxygenase and kynurenine formamidase, respectively (Fig. 2a). These two genes are present in the biosynthetic gene cluster (BGC) of actinomycin D in *S. anulatus*^26^ (Supplementary Fig. 2). The amplified *acmG* and *acmF* genes from *S. anulatus* genomic DNA were cloned in pCDFDuet-1, and the resulting plasmid pKN1 was transformed into *E. coli* BAP1 strain. The resulting *E. coli* strain with pKN1 was able to produce L-kynurenine in detectable amounts (i.e., 54.35 ± 6.92 mg/L) in R/2 medium supplemented with 5 mM L-tryptophan (Fig. 2a); all the subsequent kynurenine construction experiments were conducted under this condition unless stated otherwise. However, the presence of *N*-formyl-L-kynurenine, a precursor of L-kynurenine, was also detected according to the LC-MS analysis (Supplementary Fig. 3a,b). Detection of *N*-formyl-L-kynurenine suggests the suboptimal expression of *acmF*, and therefore, N-terminus His_6_-tagged AcmF (resulting in the plasmid pKN2) was examined for the L-kynurenine biosynthesis. As a result, the use of pKN2 generated greater titer of L-kynurenine (i.e., 87.61 ± 4.98 mg/L) without the detection of *N*-formyl-L-kynurenine (Fig. 2a and Supplementary Fig. 3c). N-terminus His_6_-tagged AcmG (pKN3) was also examined, and there was a slight increase in the production titer of L-kynurenine, 88.63 ± 1.01 mg/L (Fig. 2a).

**Fig. 2.**
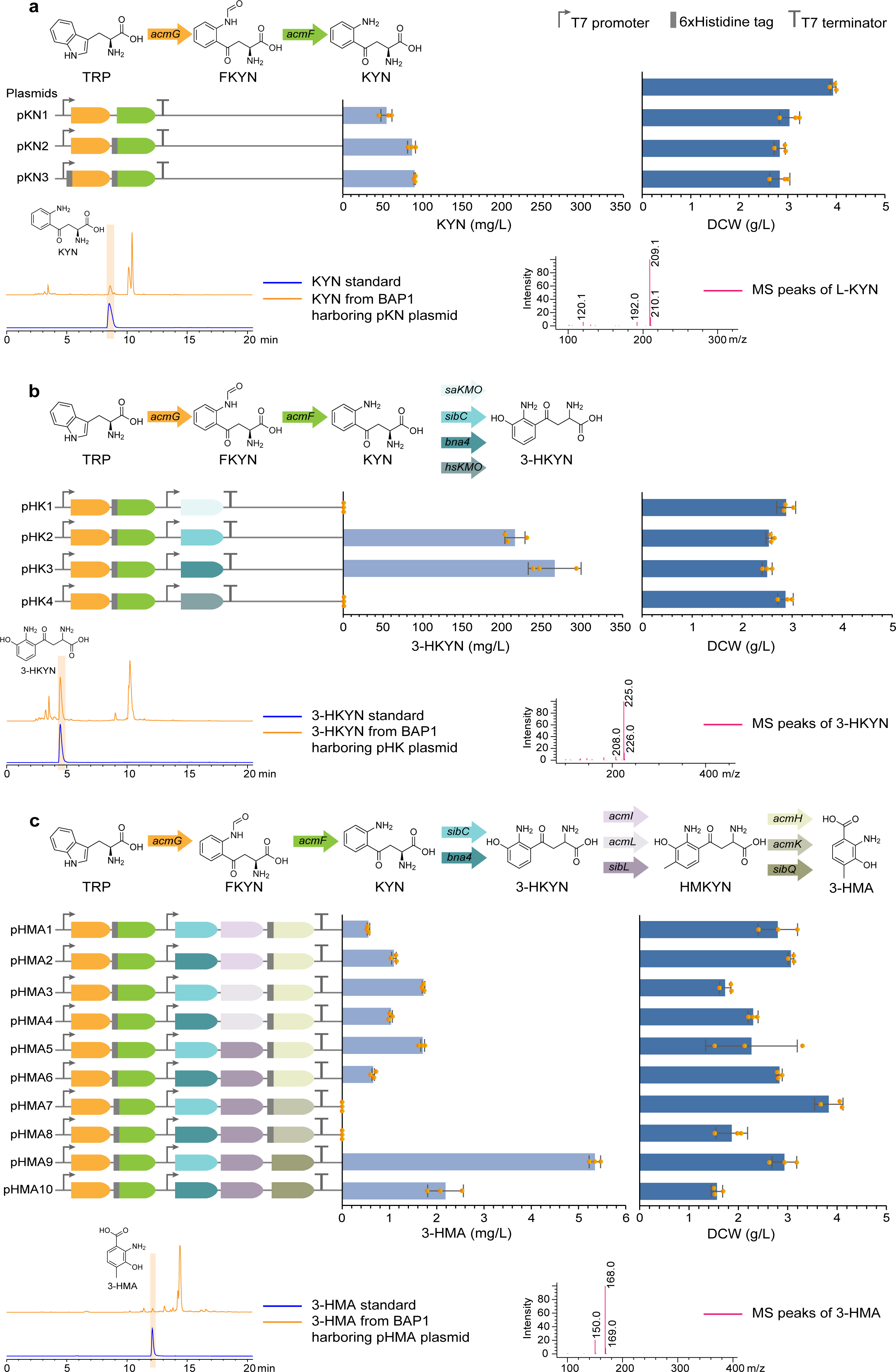
Heterologous expression of kynurenine pathway for the biosynthesis of 3-HMA. a, Production of L-kynurenine (KYN). (Upper) *acmG* and *acmF* encode tryptophan 2,3-dioxygenase and kynurenine formamidase, respectively, and they were obtained from *Streptomyces anulatus*. (Middle) KYN titer and dry cell weight (DCW) for *E. coli* BAP1 strains harboring three different pKN plasmids. (Lower) HPLC (left) and LC-MS (right) data of KYN. **b,** Production of 3-hydroxykynurenine (3-HKYN). (Upper) *saKMO*, *sibC*, *bna4*, and *hsKMO* all encode kynurenine 3-monooxygenase, which were derived from *S. anulatus*, *S. sibiricum, S. cerevisiae* and *H. sapiens*, respectively. (Middle) 3-HKYN titer and DCW for *E. coli* BAP1 strains harboring four different pHK plasmids. (Lower) HPLC (left) and LC-MS (right) data of 3-HKYN. **c,** Production of 3-hydroxy-4-methyl anthranilic acid (3-HMA). (Upper) *acmI, acmL* and *sibL* all encode methyltransferase; *acmI* and *acmL* were obtained from *S. anulatus*, and *sibL* was obtained from *S. sibiricum*. Likewise, *acmH*, *acmK* and *sibQ* all encode kynureninase; *acmH* and *acmK* were obtained from *S. anulatus*, and *sibQ* was obtained from *S. sibiricum*. (Middle) 3-HMA titer and DCW for *E. coli* BAP1 strains harboring ten different pHMA plasmids. (Lower) HPLC (left) and LC-MS (right) data of 3-HMA. (**a-c**) All the strains were grown in R/2 medium supplemented with 5 mM L-tryptophan (TRP) and appropriate antibiotics using 250 mL baffled flasks. Data present mean values and their standard deviation from triplicate experiments of flask cultivation. Metabolite abbreviations: FKYN, *N*-formyl-L-kynurenine; and HMKYN, 3-hydroxy-4-methylkynurenine.

After confirming the biosynthesis of L-kynurenine in *E. coli*, further genes encoding kynurenine 3-monooxygenase (KMO) were introduced to produce the subsequent intermediate 3-hydroxykynurenine. First, the speculated KMO-encoding gene from *S. anulatus* (*saKMO* in Fig. 2b) was introduced in the pKN2 plasmid; this gene was expected to encode KMO according to a previous study^27^. The resulting pHK1 plasmid was transformed into *E. coli*, but the resulting strain could not produce any 3-hydroxykynurenine (Fig. 2b). Therefore, three additional KMO-encoding genes were explored, including: *sibC* from *S. sibiricum*^19^; *bna4* from *S. cerevisiae*^28^ and *hsKMO* from *H. sapiens*^29^, which are all known to biosynthesize 3-hydroxykynurenine. The resulting respective plasmids pHK2, pHK3 and pHK4, all having the codon-optimized sequences (Supplementary Table 4), were individually transformed into the *E. coli* BAP1 strain. Among the three genes, *sibC* from *S. sibiricum* and *bna4* from *S. cerevisiae* successfully led to the biosynthesis of 3-hydroxykynurenine at 215.09 ± 33.04 mg/L and 264.85 ± 12.57 mg/L, respectively (Fig. 2b).

Final target metabolite to produce from kynurenine pathway was 3-HMA, which requires two additional reactions from 3-hydroxykynurenine that are catalyzed by methyltransferase and kynureninase (Fig. 2c). Here, because the standard for 3-hydroxy-4-methyl-L-kynurenine (HMKYN), a metabolite after 3-hydroxykynurenine, was not available for the detection, two genes for methyltransferase and kynureninase were incorporated into a plasmid together for the biosynthesis of 3-HMA. For the optimal biosynthesis of 3-HMA, three homologous genes for methyltransferase and another three homologous genes for kynureninase were examined in combination together with *sibC* or *bna4*. A particular emphasis was put on the genes from *S. anulatus* and *S. sibiricum*, and a total of ten combinations, each consisting of three genes, were examined as a result (Fig. 2c and Supplementary Tables 1-4). Among the corresponding ten different plasmids, the *E. coli* strain with pHMA9 produced the greatest titer of 3-HMA, 5.35 ± 0.12 mg/L. The plasmid pHMA9 contains following five genes: *acmG* and *acmF* from *S. anulatus*; and *sibC*, *sibL* and *sibQ* from *S. sibiricum*. This 3-HMA titer was substantially greater than the titers from the strains harboring pHMA10 (*bna4* in place of *sibC* in comparison with pHMA9), pHMA5 (*acmH* in place of *sibQ*), and pHMA3 (*acmL* in place of *sibL*, and *acmH* in place of *sibQ*), which generated 2.18 ± 0.37 mg/L, 1.70 ± 0.039 mg/L and 1.72 ± 0.02 mg/L of 3-HMA, respectively. It should be noted that the last three genes in kynurenine pathway that encode KMO, methyltransferase and kynureninase were all obtained from *S. sibiricum*, and this combination showed the best catalytic performance for converting 3-hydroxykynurenine to 3-HMA. Also, at this stage for the 3-HMA biosynthesis, 10 mM L-methionine^30^ was added to the R/2 medium along with 5 mM L-tryptophan because of the methylation reaction by methyltransferase (encoded by *sibL*). Based on its best production performance, the plasmid pHMA9 was used in the subsequent strains to be developed.

### Enhanced production of 3-HMA

Upon the heterologous expression of kynurenine pathway in *E. coli* producing 3-HMA, three metabolic engineering strategies were implemented to enhance the 3-HMA production, including inhibition of a competing pathway and optimal intracellular supply of SAM and PLP that are needed as cofactors for the last two biosynthetic steps of 3-HMA (Fig. 3). First, *aspC* gene encoding aspartate aminotransferase in the strain BAP1 with pHMA9 was knocked out to concentrate more fluxes toward 3-HMA through 3-hydroxykynurenine (Figs. 1 and 3). Motivation behind this strategy stems from a previous study reporting that aspartate aminotransferase in *E. coli* is capable of converting L-kynurenine to kynurenic acid^31^ in addition to its main roles in the biosynthesis of L-aspartate, L-phenylalanine and L-tyrosine, and other metabolites through transamination^32^. To confirm this additional activity of aspartate aminotransferase, the BAP1 strain without any plasmid was cultured in the R/2 medium with addition of L-kynurenine, and as a result, production of kynurenic acid was detected (Supplementary Fig. 4a). Kynurenic acid was also detected as a byproduct in the culture medium along with 3-HMA from *E. coli* BAP1 with pHMA9 (Supplementary Fig. 4b). Therefore, *aspC* gene was knocked out to inhibit the metabolic pathway that converts L-kynurenine to kynurenic acid, diverting fluxes away from 3-hydroxykynurenine. As a result of the *aspC* knockout, 15.35 ± 0.11 mg/L of 3-HMA was produced from the BAP1 Δ*aspC* with pHMA9, which is approximately three times greater than the parental strain’s (BAP1 with pHMA9) production titer. Also, kynurenic acid was substantially reduced from 32.41 ± 1.38 mg/L to 7.98 ± 0.23 mg/L (Fig. 3a and Supplementary Fig. 4c).

**Fig. 3.**
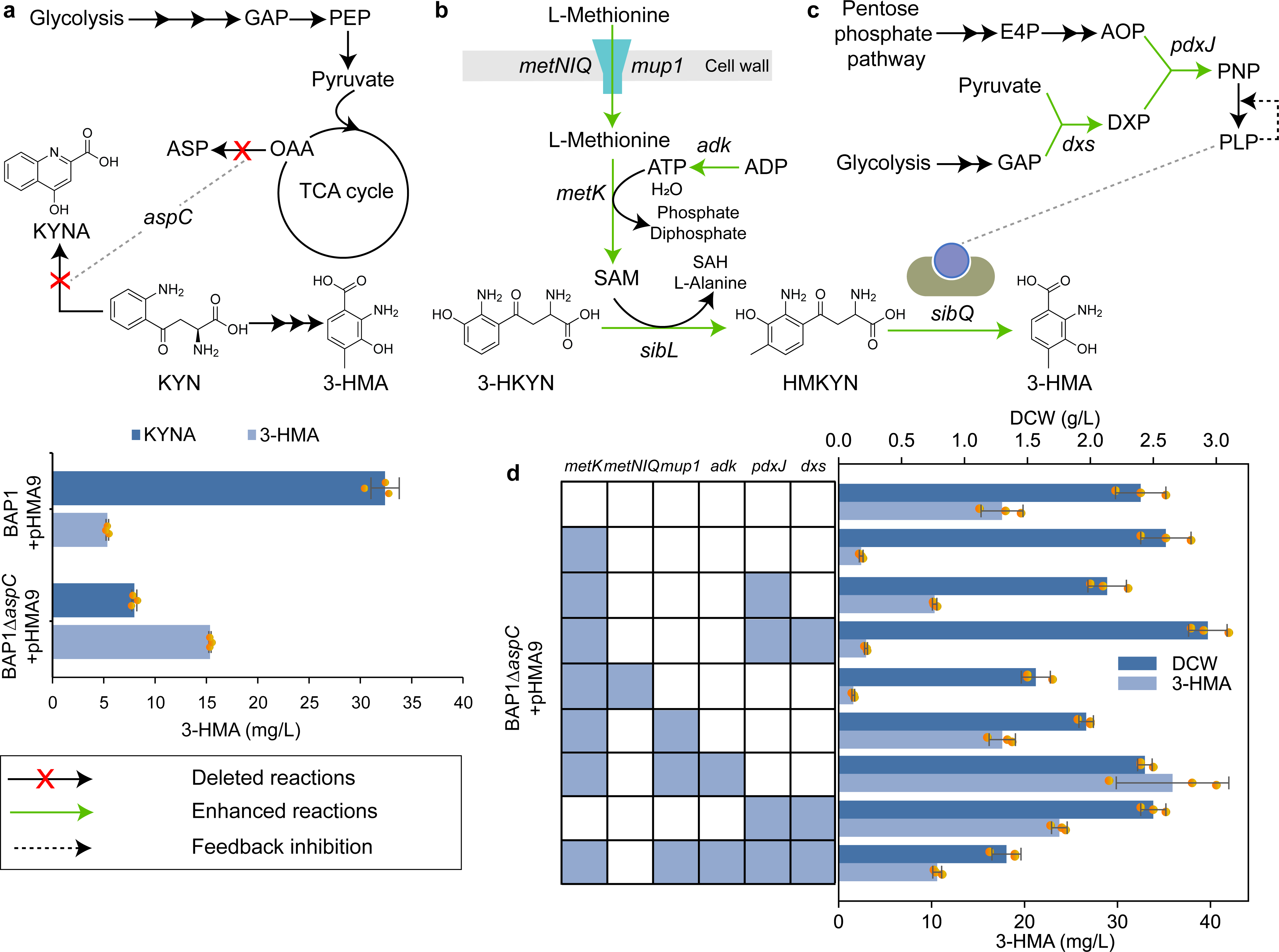
Metabolic engineering strategies for the enhanced production of 3-HMA. **a**, (Upper) Knockout of *aspC* to remove the competing pathway, and to direct fluxes toward 3-HMA. (Lower) Titers of kynurenic acid (KYNA) and 3-HMA before and after the *aspC* knockout. Aspartate aminotransferase (encoded by *aspC*) can convert oxaloacetate (OAA) to L-aspartate (ASP), and also L-kynurenine (KYN) to KYNA. **b,** Overexpression of *metK*, *metNIQ*, *mup1*, and/or *adk* to enhance the SAM biosynthesis, thereby increasing the activity of methyltransferase (*sibL*). **c,** Overexpression of *dxs* and/or *pdxJ* to enhance the pyridoxal 5’-phosphate (PLP) biosynthesis, thereby optimizing the activity of kynureninase (*sibQ*). **d,** Dry cell weight (DCW) and 3-HMA titer as a result of overexpressing the genes mentioned above (b and c). Data present mean values and their standard deviation from triplicate experiments of flask cultivation. Metabolite abbreviations: 3-HKYN, 3-hydroxykynurenine; AOP, 3-amino-2-oxopropyl phosphate; DXP, deoxyxylulose 5-phosphate; E4P, erythrose 4-phosphate; GAP, glyceraldehyde 3-phosphate; HMKYN, 3-hydroxy-4-methylkynurenine; KYN, L-kynurenine; PEP, phosphoenolpyruvate; PNP, pyridoxine 5’-phosphate; SAH, *S*-adenosyl-L-homocysteine; and SAM, *S*-adenosyl-L-methionine.

Next, the conversion of 3-HKYN to HMKYN by *sibL*-encoded methyltransferase requires *S*-adenosyl-L-methionine (SAM) that provides a methyl group for this reaction^20^ (Fig. 3b). Therefore, to further enhance the 3-HMA titer, optimizing the intracellular supply of SAM was selected as a next engineering target. For this, *S*-adenosylmethionine synthase encoded by *metK*^34^ was overexpressed (using the pET-30a-based plasmid pSAM) to improve the SAM availability. However, the resulting BAP1Δ*aspC* strain with pHMA9 and pSAM could only produce 2.42 ± 0.20 mg/L of 3-HMA (Fig. 3d).

Overexpression of *metK* was attempted again, but this time, with additional genes, including those involved in PLP biosynthesis (Fig. 3c,d). The final step of the 3-HMA biosynthesis is catalyzed by *sibQ*-encoded kynureninase^20^, converting HMKYN to 3-HMA, and it requires PLP as a cofactor. In *E. coli*, both glycolysis and pentose phosphate pathway can generate precursors that are needed for the PLP biosynthesis^35–37^ (Fig. 3c). Thus, the native *pdxJ* gene encoding pyridoxine 5’-phosphate synthase was overexpressed along with *metK* by using the resulting pSP1 vector (obtained by integrating *pdxJ* in pSAM vector; Supplementary Tables 1 and 2). As a result, the 3-HMA titer improved to 10.32 ± 0.24 mg/L in the strain BAP1Δ*aspC* with pHMA9 and pSP1 (Fig. 3d). However, the titer was still less than 17.58 ± 2.25 mg/L of 3-HMA from the BAP1Δ*aspC* strain with only pHMA9. In an attempt to enhance the titer further by increasing PLP availability, *dxs* gene encoding 1-deoxy-D-xylulose-5-phosphate synthase was overexpressed together with *metK* and *pdxJ* using vector pSP2 (obtained by integrating *dxs* in pSP1; Supplementary Tables 1 and 2). This attempt improved the growth of the resulting strain BAP1Δ*aspC* with pHMA9 and pSP2, but the 3-HMA titer decreased to 2.95 ± 0.16 mg/L (Fig. 3d).

At this stage, optimization process of the SAM and PLP biosynthesis was implemented separately. For the SAM biosynthesis, the native *metNIQ* cassette^38^ was overexpressed using pET-30a-based plasmid pSAM1 that regulates L-methionine import along with *metK* in the BAP1Δ*aspC* with pHMA9. However, the 3-HMA titer was rather substantially decreased as compared to the strain BAP1Δ*aspC* with pHMA9 (Fig. 3d). As the second attempt, another methionine permease (encoded by *mup1*) from *S. cerevisiae* was explored as there was report on the expression of *mup1* in a heterologous host *Pichia pastoris*, which led to the enhanced SAM production^39^. Hence, along with overexpression of the native *metK* gene, codon-optimized *mup1* gene was also expressed in *E. coli* using pET-30a as the expression vector (pSAM2). This time, 3-HMA titer was successfully increased to 17.60 ± 1.41 mg/L. Also, because SAM biosynthesis requires a mole of ATP, the native *adk* gene encoding adenylate kinase was overexpressed along with *metK* and *mup1* (pSAM3) for further enhanced 3-HMA titer. Indeed, the resulting strain BAP1Δ*aspC* harboring pHMA9 and pSAM3 produced a substantially greater titer of 3-HMA, reaching 35.91 ± 6.05 mg/L from flask cultivation (Fig. 3d). For enhancing the PLP biosynthesis, overexpression of *pdxJ* and *dxs* genes was attempted using pRSFDuet-A-based pPLP. The resulting strain BAP1Δ*aspC* with pHMA9 and pPLP, overexpressing *pdxJ* and *dxs*, produced 23.74 ± 0.84 mg/L of 3-HMA, which was lower than the titer from the BAP1Δ*aspC* strain with pHMA9 and pSAM3, overexpressing only genes associated with the SAM biosynthesis (Fig. 3d).

Finally, in the BAP1Δ*aspC* strain with pHMA9, the two plasmids, pSAM3 and pPLP1, were co-expressed. These two plasmids had shown the two best production titers of 3-HMA when independently expressed. However, the resulting strain showed poorer growth, and 3-HMA titer also dropped significantly to 10.60 ± 0.47 mg/L (Fig. 3d). As a result, the BAP1Δ*aspC* strain with pHMA9 and pSAM3 was used for further engineering as it so far showed the best 3-HMA production titer (35.91 ± 6.05 mg/L) through the optimal SAM biosynthesis.

### Production of actinocin from 3-HMA

To explore the potential of kynurenine pathway for the production of medicinal molecules, we attempted to produce actinocin as a primary target molecule. Actinocin is a chromophore of actinomycin D, and is formed via condensation of two 3-HMA molecules. This condensation reaction has been reported to be catalyzed by a spore pigment-synthesizing laccase (encoded by *cotA*) of *Bacillus subtilis*^40, 41^. To express laccase in *E. coli*, its *cotA* gene was amplified from the genomic DNA of *B. subtilis* 168, and cloned in pRSFDuet-A vector, resulting in pACN1 plasmid. Because the standard for actinocin was not available, the successful expression of laccase was first confirmed using SDS-PAGE for the BAP1Δ*aspC* strain with pACN1 (Fig. 4a). To further confirm the conversion of 3-HMA to actinocin, 3-HMA was supplemented to the BAP1Δ*aspC* strain with pACN1. HPLC analysis showed that the strain generated HPLC peaks potentially indicative of actinocin production (Fig. 4b). The BAP1Δ*aspC* strain was subsequently transformed with pHMA9, pSAM3 and pACN1, each responsible for the biosynthesis of 3-HMA, SAM and actinocin, respectively. The resulting strain produced 8.72 ± 0.39 mg/L of actinocin (Fig. 4c). The produced actinocin was purified using fraction collector and confirmed by NMR (Supplementary Fig. 5 and Methods).

**Fig. 4.**
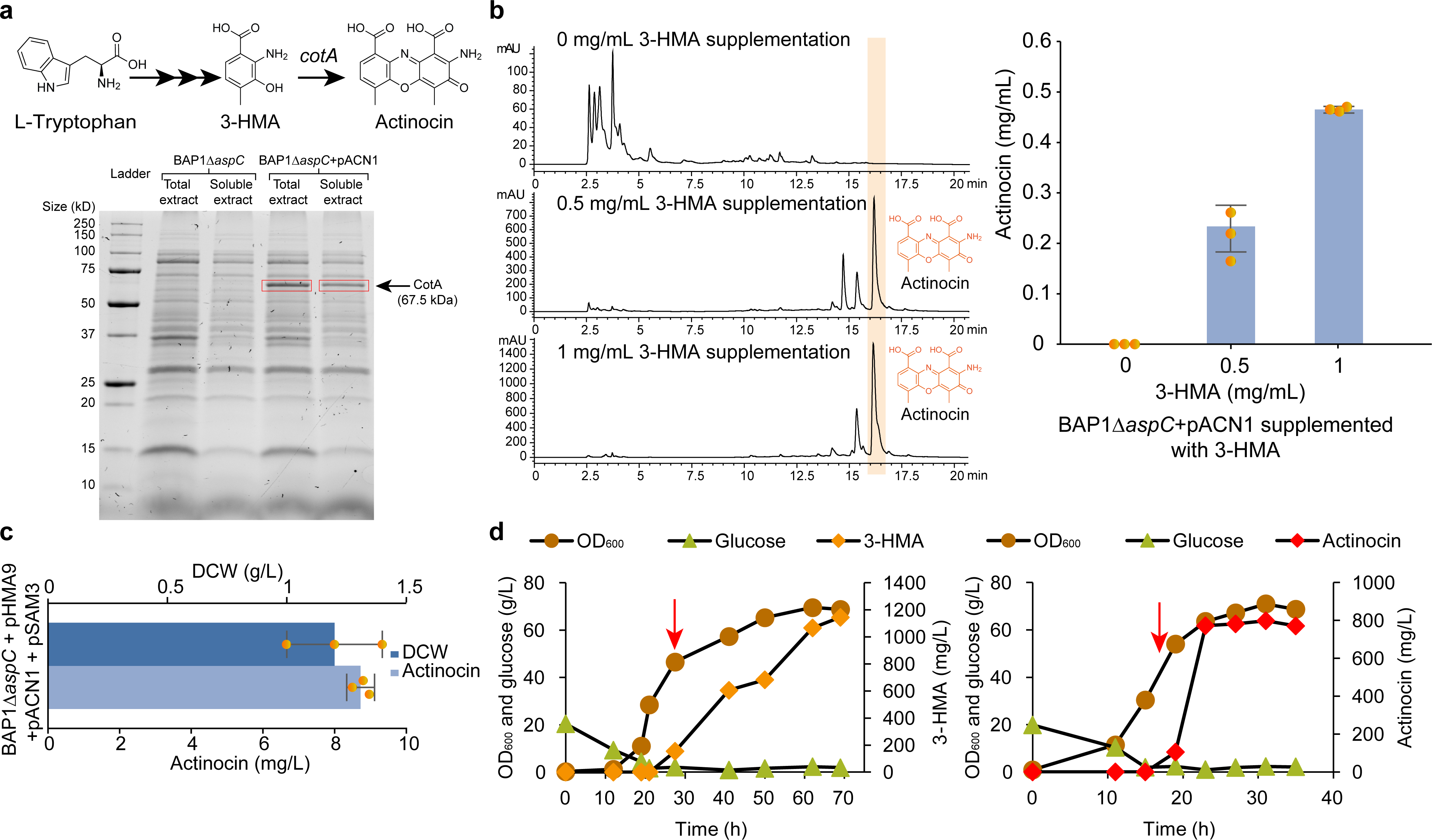
Production of actinocin from 3-HMA. **a**, (Upper) Production of actinocin by additionally expressing *cotA* encoding laccase. (Lower) SDS-PAGE results confirming the successful expression of laccase in *E. coli* BAP1Δ*aspC* strain harboring pACN1. **b,** 3-HMA feeding experiment to confirm the successful production of actinocin via laccase expressed in the BAP1Δ*aspC* strain with pACN1. (Left) Detection of HPLC peaks potentially corresponding to actinocin as a result of 3-HMA supplementation. (Right) Actinocin titers as a result of 3-HMA supplementation. **c,** Dry cell weight (DCW) and actinocin titer from flask cultivation. **d,** Fed-batch fermentation profiles of BAP1Δ*aspC* with pHMA9 and pSAM3 for producing 3-HMA production (left), and BAP1Δ*aspC* with pHMA9, pSAM3 and pACN1 for producing actinocin (right). Red arrow indicates the point of isopropyl β-D-1-thiogalactopyranoside (IPTG) induction. (**b** and **c**) Flask cultivations were performed in R/2 medium supplemented with 5 mM L-tryptophan, 10 mM L-methionine, and appropriate antibiotics. Data present mean values and their standard deviation from triplicate experiments.

Finally, the two strains were subjected to fed-batch fermentation: BAP1Δ*aspC* with pHMA9 and pSAM3, producing 3-HMA, as a reference, and BAP1Δ*aspC* with pHMA9, pSAM3 and pACN1 for the production of actinocin (Fig. 4d). As a result, BAP1Δ*aspC* with pHMA9 and pSAM3 produced 1145 mg/L of 3-HMA (left data in Fig. 4d), and BAP1Δ*aspC* strain with pHMA9, pSAM3 and pACN1 produced 799 mg/L of actinocin (right data in Fig. 4d).

### De novo production of actinocin from glucose

Further metabolic engineering was conducted to produce actinocin from glucose without supplementation of exogenous L-tryptophan. Such de novo production would require the enhanced biosynthesis of L-tryptophan, which, however, is subject to feedback inhibition. To address this issue, four genes were engineered. First, the engineered *aroG* and *trpE* genes, encoding feedback-resistant 3-deoxy-D-arabino-heptulosonate 7-phosphate (DAHP) synthase (AroG^A146N^) and feedback-resistant anthranilate synthase (TrpE^S40F^), respectively, were targeted for overexpression on the basis of a previous study^42^. These two enzymes are feedback-resistant to L-phenylalanine^43^ and L-tryptophan^44^, respectively, and they were considered crucial for the enhanced biosynthesis of L-tryptophan. Second, *aroL* gene encoding shikimate kinase II overexpression is also known to be beneficial for L-tryptophan biosynthesis^45^. The above three genes were incorporated in the pTac15K plasmid for co-overexpression, resulting in pTRP plasmid^42^. Finally, *tnaA* gene encoding tryptophanase was knocked out, generating the strain BAP1Δ*aspC*Δ*tnaA*. Tryptophanase is responsible for conversion of L-tryptophan to indole in *E. coli*^46, 47^. As a result, 3-HMA-producting strain harboring pHMA9, pSAM4 and pTRP was constructed (upper scheme in Fig. 5a). It should be noted that pSAM4 was constructed from pSAM3 by replacing kanamycin with ampicillin because pTRP, originally designed by Du et al. (2018), already contained kanamycin as a resistance marker. The resulting strain successfully produced 10.68 ± 1.79 mg/L of 3-HMA in flask cultivation without any L-tryptophan supplementation (Fig. 5b). For the production of actinocin from glucose, pSAM3 and pACN1 were integrated as pACN2 to reduce the number of plasmids (lower scheme in Fig. 5a). The resulting strain BAP1Δ*aspC*Δ*tnaA* with pHMA9, pACN2 and pTRP produced 2.76 ± 0.13 mg/L of actinocin directly from glucose (Fig. 5b).

**Fig. 5.**
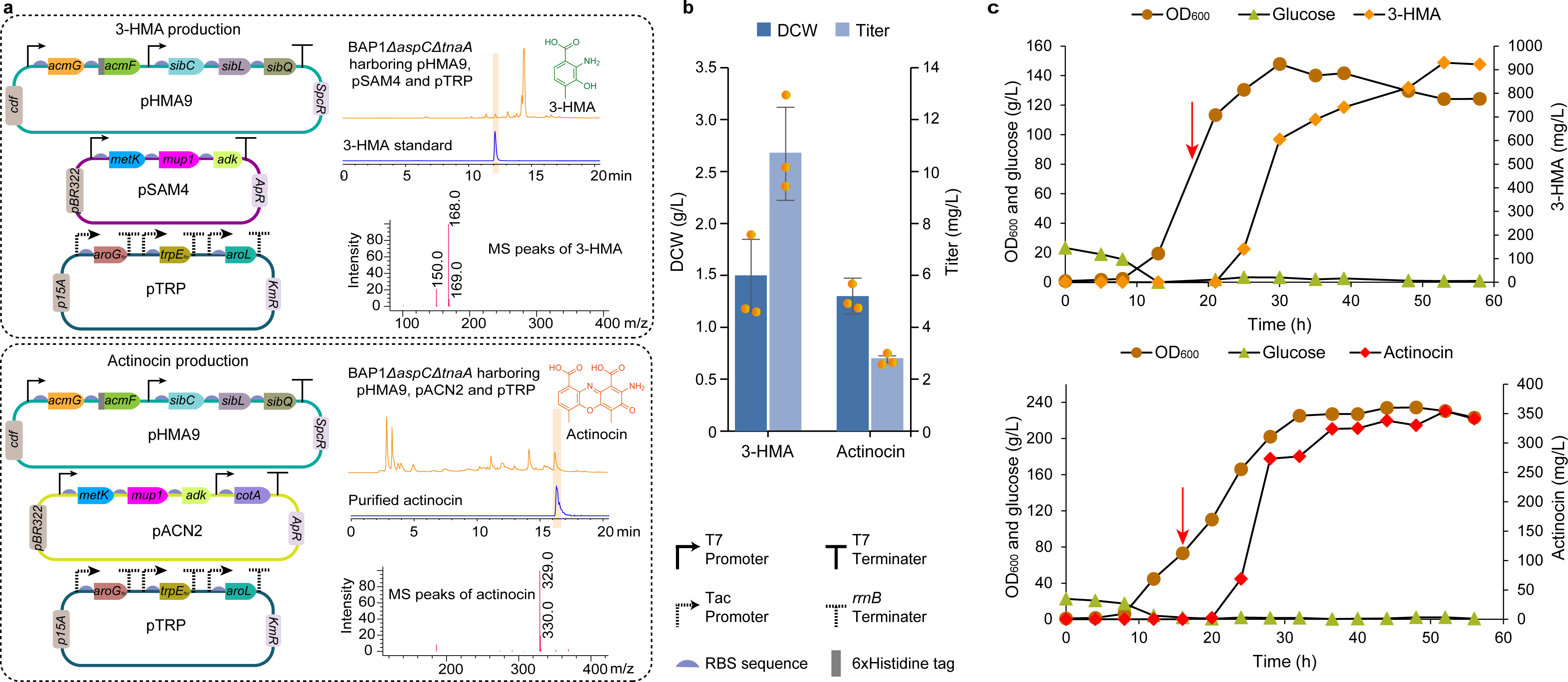
De novo production of 3-HMA and actinocin from glucose. **a**, Plasmid maps for the production of 3-HMA (upper) and actinocin (lower). HPLC (top) and LC-MS (bottom) data of 3-HMA and actinocin are also presented next to the plasmid maps. **b,** Dry cell weight (DCW) and titers for 3-HMA and actinocin. Data present mean values and their standard deviation from triplicate experiments of flask cultivation. **c,** Fed-batch fermentation profiles of BAP1Δ*aspC*Δ*tnaA* with pHMA9, pSAM4 and pTRP for producing 3-HMA (upper), and BAP1Δ*aspC*Δ*tnaA* with pHMA9, pACN2 and pTRP for producing actinocin (lower). Red arrow indicates the point of isopropyl β-D-1-thiogalactopyranoside (IPTG) induction.

The two strains, producing 3-HMA and actinocin from glucose, were subjected to fed-batch fermentation using a 6.6 L bioreactor to properly examine their production performances. As a result, 929 mg/L of 3-HMA (upper data in Fig. 5c) and 354 mg/L of actinocin (lower data in Fig. 5c) were produced from these two strains. These production titers were greater than those previously reported for other kynurenine pathway-derived molecules, including kynurenic acid from *Yarrowia lipolytica*^48^ (17.7 mg/L from 150 h of cultivation with precursor supplementation) and cinnabarinic acid from *Pseudomonas chlororaphis*^49^ (136.2 mg/L from fed-batch fermentation using shake flasks) (Supplementary Table 5).

### Enhanced production of actinocin based on metabolome analysis

The actinocin titer from the fed-batch fermentation by using glucose was 354 mg/L (lower data in Fig. 5c), which was much lower than the titer from the fed-batch fermentation with supplementation of L-tryptophan (799 mg/L; right data in Fig. 4d). Thus, metabolic engineering was further performed on the basis of metabolome analysis. It was thought that metabolome analysis would reveal metabolites with particularly high concentration in the 3-HMA-producing strain (BAP1Δ*aspC*Δ*tnaA* with pHMA9, pSAM4 and pTRP) in comparison with the control strains (*E. coli* BAP1 and L-kynurenine-producing strain *E. coli* BAP1 with pKN2; Fig. 6). Identification of such metabolites would reveal enzymes that are not efficiently catalyzing the reaction, thereby suggesting next overexpression targets. In this study, 353 metabolites in the abovementioned three strains were subjected to absolute quantification (Fig. 6, Supplementary Data 1, and Methods). The resulting metabolome profiles showed that fructose 1,6-bisphosphate and isocitrate appeared to be substantially accumulated in the 3-HMA-producing strain, compared with those in the control strains (Fig. 6). There were additional metabolites substantially accumulated in the 3-HMA-producing strain, including SAM, ATP, 3-dehydroshikimate (DHS), shikimate (SHI), shikimate 3-phosphate (S3P), and anthranilate (ANT). The increased concentration of these metabolites was likely attributed to the presence of pSAM4 for SAM and ATP, and pTRP for DHS, SHI, S3P, and ANT in the 3-HMA-producing strain (Figs. 1 and 6). In this study, we focused on metabolites in glycolysis and TCA cycle that can provide global metabolic effects. Consequently, fructose 1,6-bisphosphate aldolase (encoded by *fbaA*) and isocitrate dehydrogenase (encoded by *icd*) were chosen as overexpression targets as they are responsible for the generation of fructose 1,6-bisphosphate and isocitrate, respectively.

**Fig. 6.**
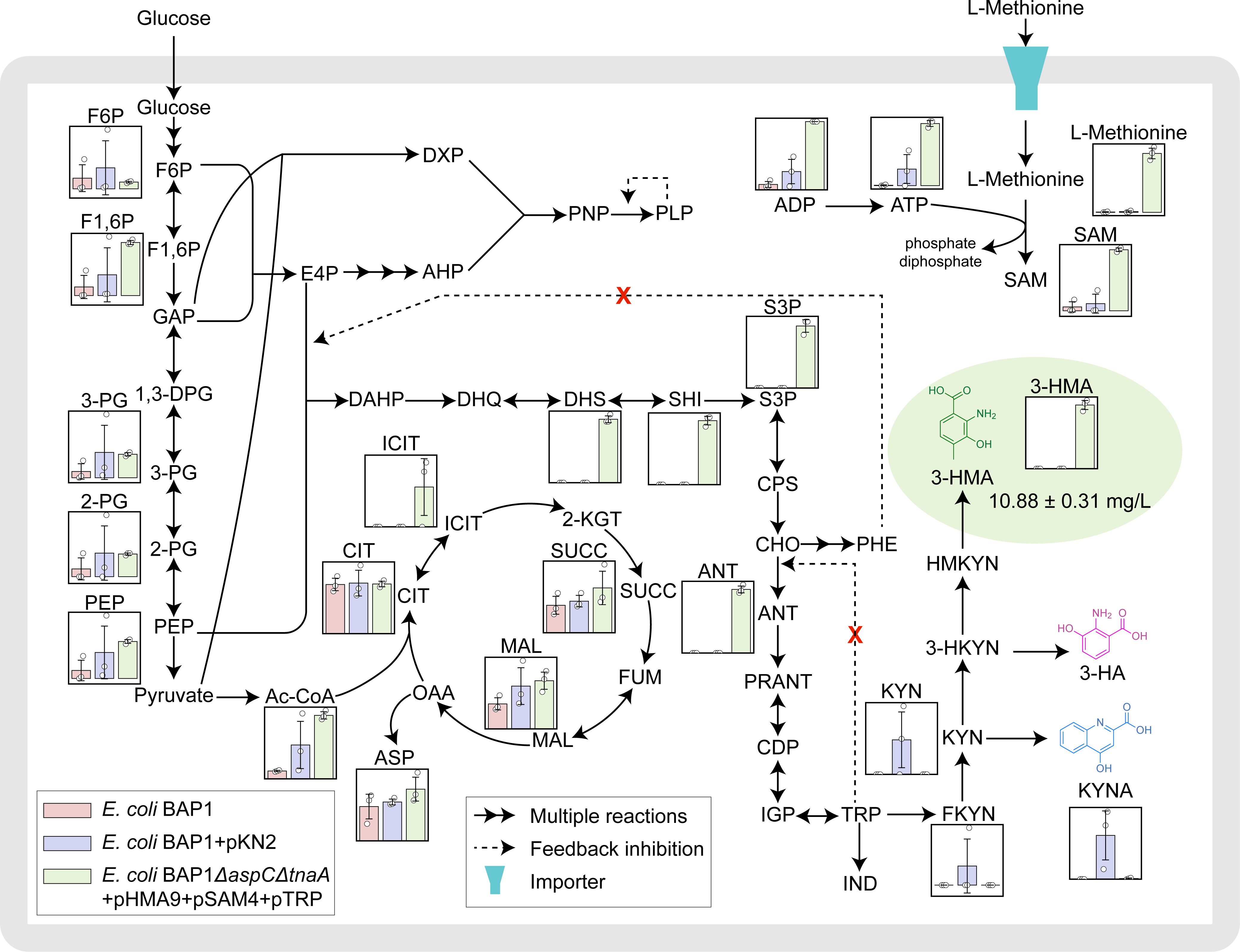
Metabolome analysis to identify additional gene manipulation targets. Metabolome profiles of the *E. coli* BAP1 strain, *E. coli* BAP1 with pKN2 producing L-kynurenine with L-tryptophan supplementation, and the final engineered strain *E. coli* BAP1Δ*aspC*Δ*tnaA* with pHMA9, pSAM4 and pTRP producing 3-HMA from glucose. Metabolites were subjected to absolute quantification, and their concentrations are presented as pmol/OD mL. Metabolite abbreviations: 1,3-DPG, 1,3-diphospho D-glycerate; 2-PG, 2-phosphoglycerate; 3-HA, 3-hydroxyanthranilic acid; 3-HKYN, 3-hydroxykynurenine; 3-HMA, 3-hydroxy-4-methyl anthranilic acid; 3-PG, 3-phosphoglycerate; Ac-CoA, acetyl-Coenzyme A; ANT, anthranilate; ASP, aspartate; AOP, 3-amino-2-oxopropyl phosphate; CDP, 1-(2-carboxyphenylamino)-1-deoxy-D-ribulose-5-phosphate; CHO, chorismate; CIT, citrate; CPS, 5-*o*-(1-carboxyvinyl)-3-phosphoshikimate; DAHP, 3-deoxy-arabino-heptulonate-7-phosphate; DHQ, 3-dehydroquinate; DHS, 3-dehydroshikimate; DXP, deoxyxylulose 5-phosphate; E4P, erythrose 4-phosphate; F1,6P, fructose 1,6-bisphosphate; F6P, fructose 6-phosphate; FKYN, *N*-formyl-L-kynurenine; FUM, fumarate; GAP, glyceraldehyde 3-phosphate; HMKYN, 3-hydroxy-4-methylkynurenine; ICIT, isocitrate; IGP, indole 3-glycerol phosphate; IND, indole; KGT, α-ketoglutarate; KYN, L-kynurenine; MAL, malate; OAA, oxaloacetate; PEP, phosphoenolpyruvate; PLP, pyridoxal 5’-phosphate; PNP, pyridoxine 5’-phosphate; PRANT, *N*-(5-phospho-D-ribosyl) anthranilate; S3P shikimate-3-phosphate; SAH, *S*-adenosyl-L-homocysteine; SAM, *S*-adenosyl-L-methionine; SUCC, succinate; SHI, shikimate; and TRP, L-tryptophan.

Individual overexpression of *icd* (using plasmid pIcd) and *fbaA* (using plasmid pFbaA), and their co-overexpression (using plasmid pIF) were conducted using the 3-HMA-producing strain (BAP1Δ*aspC*Δ*tnaA* with pHMA9, pSAM4 and pTRP1), and also the actinocin-producing strain (BAP1Δ*aspC*Δ*tnaA* with pHMA9, pACN2 and pTRP1) (Fig. 7a,b); pTRP1 is a modified version of pTRP where positions of the genes were rearranged to avoid the gene loss (e.g., engineered *trpE*) that actually took place during cultivations. As a result of the additional expression studies, co-overexpression of *icd* and *fbaA* led to the best production of 3-HMA (left data in Fig. 7a) and actinocin (right data in Fig. 7a) from flask cultivation. BAP1Δ*aspC*Δ*tnaA* with pHMA9, pSAM4, pTRP1 and pIF produced 15.04 ± 0.16 mg/L of 3-HMA, and BAP1Δ*aspC*Δ*tnaA* with pHMA9, pACN2, pTRP1 and pIF produced 17.87 ± 2.16 mg/L of actinocin (Fig. 7a). Interestingly, for the first time in this study, actinocin titer appeared to be greater than 3-HMA titer from their independent cultivation experiments. Before the co-overexpression of *icd* and *fbaA*, 3-HMA titer was always greater than actinocin titer from independent cultivations: i.e., 1145 mg/L of 3-HMA versus 799 mg/L of actinocin, both with L-tryptophan supplementation (Fig. 4d); and 929 mg/L of 3-HMA versus 354 mg/L of actinocin, both from glucose (Fig. 5c). The co-expression somehow contributed to the more efficient conversion of 3-HMA to actinocin. For the actinocin-producing strain, its growth also improved as a result of the co-overexpression, in comparison with individual expression of the two genes (Fig. 7c). Finally, the final actinocin-producing strain (BAP1Δ*aspC*Δ*tnaA* with pHMA9, pACN2, pTRP1 and pIF) was subjected to fed-batch fermentation in a 6.6 L bioreactor, which produced 719 mg/L of actinocin (Fig. 7c and Supplementary Fig. 6). A sharp increase in the 3-HMA titer was observed, which was quickly followed by a decrease, coinciding with an increase in the actinocin titer upon IPTG induction. As a result, actinocin titer was recovered by more than twofold, compared with the strain without the co-overexpression, as a result of the additional engineering based on the metabolome analysis.

**Fig. 7.**
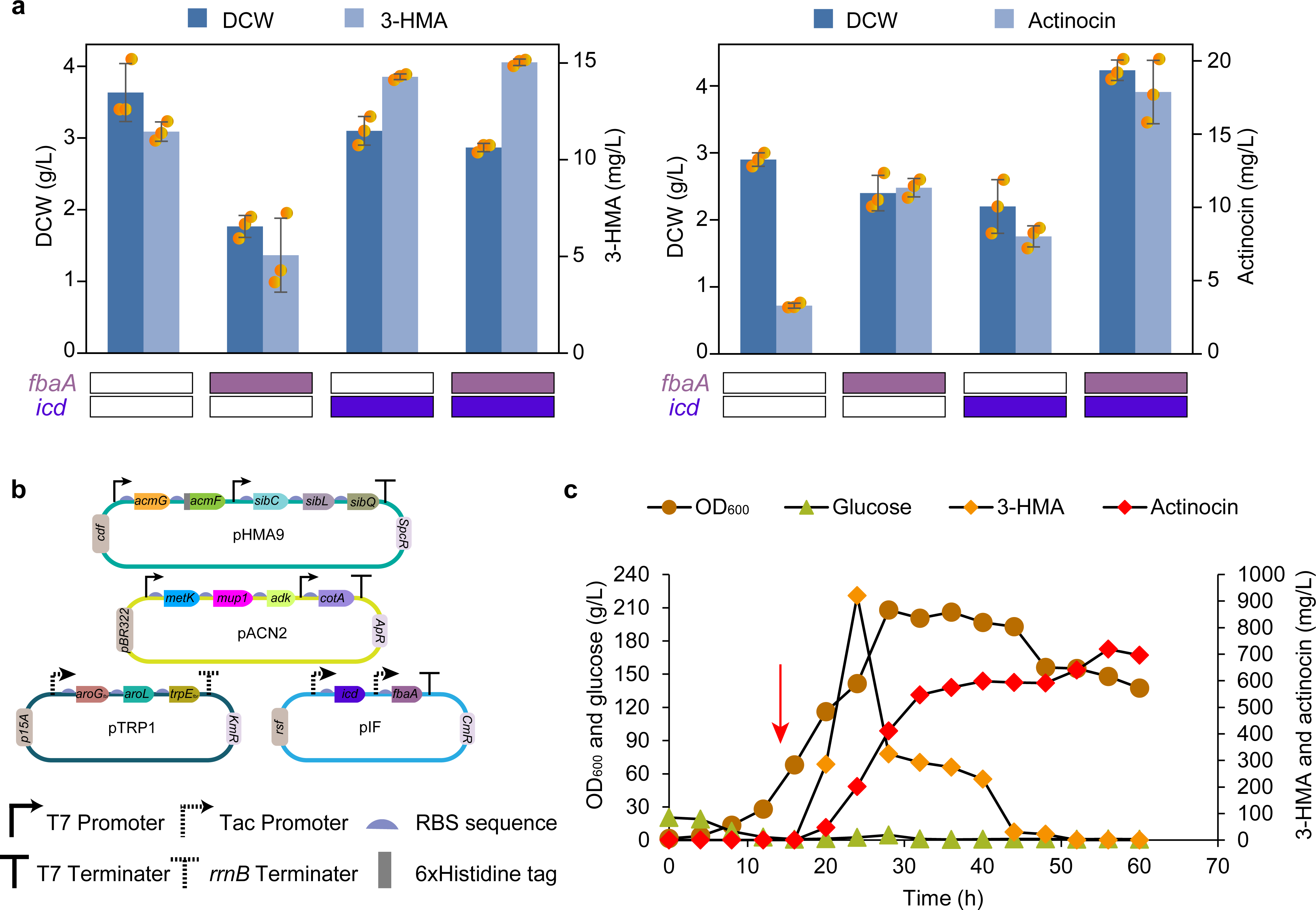
Enhanced production of actinocin based on metabolome analysis. **a**, Dry cell weight (DCW) and titers for 3-hydroxy-4-methyl anthranilic acid (3-HMA) (left) and actinocin (right) after overexpression of *fbaA* (encoding fructose-1,6-bisphosphate aldolase) and/or *icd* (encoding isocitrate dehydrogenase). These genes were expressed in the 3-HMA-producing strain (*E. coli* BAP1Δ*aspC*Δ*tnaA* with pHMA9, pSAM4 and pTRP1) and the actinocin-producing strain (*E. coli* BAP1Δ*aspC*Δ*tnaA* with pHMA9, pACN2 and pTRP1). Data present mean values and standard deviation from triplicate experiments of flask cultivation. **b,** Final plasmid map for the actinocin production using the strain *E. coli* BAP1Δ*aspC*Δ*tnaA*. For pTRP1 plasmid, it has one promoter-terminator system for the three genes, which helps avoid any possible recombination and subsequent loss of the genes. **c**, Fed-batch fermentation profile of the final engineered strain *E. coli* BAP1Δ*aspC*Δ*tnaA* with pHMA9, pACN2, pTRP1 and pIF. Red arrow indicates the point of isopropyl β-D-1-thiogalactopyranoside (IPTG) induction.

### De novo production of kynurenic acid and 3-HA from glucose

To further explore the biosynthetic potential of kynurenine pathway by using *E. coli*, we also attempted to produce two relevant kynurenine derivatives, kynurenic acid and 3-HA, from glucose as additional target molecules (Fig. 1). For kynurenic acid, the BAP1Δ*aspC*Δ*tnaA* strain was transformed with pKYNA expressing *acmG*, *acmF*, and *aspC*, and pTRP (upper scheme in Fig. 8a). The resulting strain produced 6.64 ± 0.34 mg/L of kynurenic acid solely from glucose during flask cultivation (Fig. 8b).

**Fig. 8.**
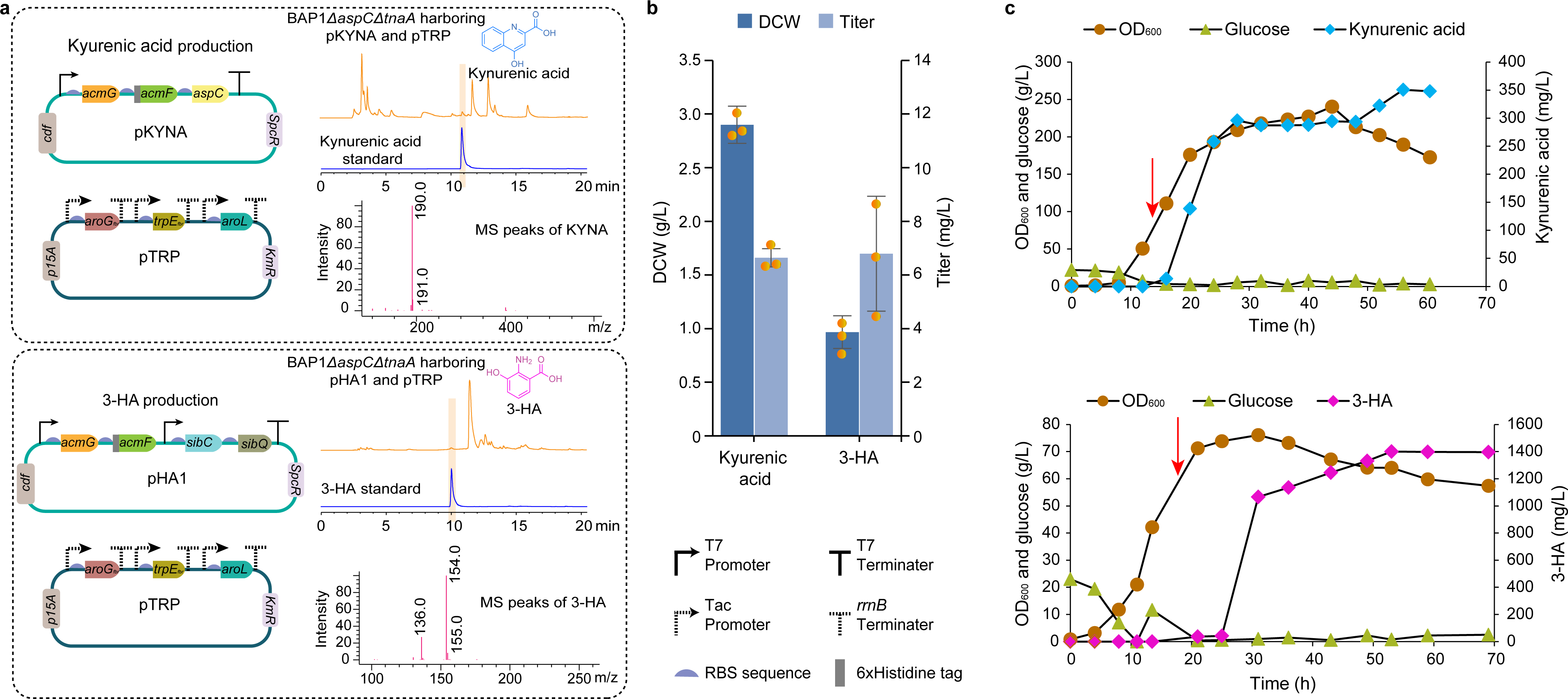
De novo production of kynurenic acid and 3-HA from glucose. **a**, Plasmid maps for the production of kynurenic acid (upper) and 3-hydroxyanthranilic acid (3-HA; lower). **b,** Dry cell weight (DCW) and titers for kynurenic acid and 3-HA. Data present mean values and their standard deviation from triplicate experiments of flask cultivation. **c,** Fed-batch fermentation profiles of BAP1Δ*aspC*Δ*tnaA* with pKYNA and pTRP for producing kynurenic acid (upper), and BAP1Δ*aspC*Δ*tnaA* with pHA1 and pTRP for producing 3-HA (lower). Red arrow indicates the point of isopropyl β-D-1-thiogalactopyranoside (IPTG) induction.

As to 3-HA, kynureninase of the kynurenine pathway has been reported to possess a broad-substrate specificity, including 3-hydroxykynurenine^20^ in addition to HMKYN discussed above (Figs. 1 and 2). Kynureninase can convert 3-hydroxykynurenine into 3-HA. Indeed, 3-HA was observed from flask cultivation of the 3-HMA-producing strains (Supplementary Fig. 7). Here, 3-HA-producing strain was constructed by using pHA1 (pCDFDuet-1 harboring *acmG*, *acmF*, *sibC* and *sibQ*) (lower scheme in Fig. 8a). The resulting strain BAP1Δ*aspC*Δ*tnaA* with pHA1 and pTRP produced 6.69 ± 2.14 mg/L of 3-HA (Fig. 8b).

Next, the two strains producing kynurenic acid and 3-HA were subjected to fed-batch fermentation using a 6.6 L bioreactor. These fed-batch fermentations led to the production of 350 mg/L of kynurenic acid (upper data in Fig. 8c) and 1401 mg/L of 3-HA (lower data in Fig. 8c), both from glucose. As with actinocin, the production titers of kynurenic acid and 3-HA were also greater than those reported by the previous studies (Supplementary Table 5). Taken together, these production studies demonstrated the use of the heterologous kynurenine pathway in *E. coli* for the efficient production of multiple medicinal molecules.

## Discussion

In this study, the kynurenine pathway was explored as it is known to be associated with multiple molecules that have potential medicinal values. For this, we have aimed at producing three medicinal molecules that can be derived from the kynurenine pathway, including actinocin, kynurenic acid and 3-HA. A common strategy behind the production of these three molecules was to first optimally express heterologous kynurenine pathway in *E. coli*. Subsequently, the strains were engineered to overproduce each molecule from glucose without supplementation of a direct precursor (i.e., L-tryptophan). Particularly, in this study, we focused on the actinocin production by enhancing the biosynthesis of its precursor 3-HMA. Systematic metabolic engineering was performed for the enhanced 3-HMA biosynthesis by examining multiple combinations of heterologous biosynthetic genes, and by optimizing the metabolic flux distribution through removal of the competing pathway, enhancement in the cofactor biosynthesis, and de novo production from glucose. Also, metabolome analysis helped identify nonintuitive, yet effective gene manipulation targets. To our knowledge, the production titers of actinocin, kynurenic acid and 3-HA from glucose appeared to be competitive, in comparison with reported studies on the kynurenine pathway-derived molecules (Supplementary Table 5).

This study also highlights several considerations that could provide future research opportunities. Kynurenine pathway starts from L-tryptophan pathway. However, kynurenine pathway seems to have lower metabolic fluxes than L-tryptophan pathway on the basis of the production titers of actinocin, kynurenic acid and 3-HA in comparison with L-tryptophan production titers previously reported^50^. Lower titers of the kynurenine pathway-derived molecules may be partly due to the nature of foreign genes expressed in a heterologous host (*E. coli* in this study). Additional optimization of the expression of both heterologous kynurenine pathway genes (Fig. 2) and native amino acid pathway fluxes (e.g., shikimate pathway; Figs. 3 and 5) may be necessary to further enhance the production titer of the kynurenine pathway-derived molecules. Utilization of omics data, for example metabolome data as demonstrated in this study, can be another considerable approach. Metabolome data in this study provided rather precise capture of a metabolic phenotype of the 3-HMA-producing strain (BAP1Δ*aspC*Δ*tnaA* with pHMA9, pSAM4, and pTRP) (Fig. 6). Metabolome data captured the increased concentration of several metabolites, which were rationally intended during the strain development, including SAM and ATP through pSAM4, and DHS, SHI, S3P, and ANT through pTRP in the 3-HMA-producing strain. Importantly, metabolome data generated nonintuitive, yet effective gene overexpression targets (Fig. 7). This study demonstrated the effective application of such omics profiling, which could additionally provide potential gene manipulation targets.

In summary, this study explored the biosynthetic potential of the kynurenine pathway. By optimizing the expression of the kynurenine pathway in *E. coli*, and metabolic flux distribution of *E. coli* metabolism, we successfully produced kynurenine pathway-derived molecules, including actinocin, kynurenic acid, and 3-HA, at competitive production level. Competitive production titers of these molecules demonstrate the potential use of kynurenine pathway as a platform for the production of other multiple medicinal molecules.

## Methods

### Bacterial strains and culture conditions

Bacterial strains used in this study are listed in Supplementary Table 1. *E. coli* DH10β (New England Biolabs) was used for plasmid construction and propagation, and *E. coli* BAP1^51^ was used as a host strain for gene expression and target compound production. *E. coli* strains were first inoculated from colonies on LB agar plates into 25 mL test tubes containing 10 mL of LB medium supplemented with appropriate antibiotics: when required, 100 μg/L of ampicillin, 50 μg/L of kanamycin, 17 μg/L of chloramphenicol, and/or 200 μg/L of spectinomycin were added to the medium (Supplementary Table 1). The test tubes were placed in a rotary shaker overnight at 37 °C and 250 rpm. One mL aliquot of each seed culture was transferred to a 250 mL baffled flask containing 25 mL of R/2 medium with 20 g/L of glucose (and 5 mM of L-tryptophan when necessary). The baffled flasks were incubated at 30°C and at 200 rpm. The R/2 medium (pH 6.8) contains the following per liter: 2 g (NH_4_)2HPO_4_, 6.75 g KH_2_PO_4_, 0.85 g citric acid, 0.8 g MgSO_4_·7H_2_O, and 5 mL trace metal solution (TMS). The TMS contains the following per liter: 0.1 M HCl, 10 g FeSO_4_·7H_2_O, 2.25 g ZnSO_4_·7H_2_O, 1 g CuSO_4_·5H_2_O, 0.58 g MnSO_4_·5H_2_O, 0.02 g Na_2_B_4_O_7_·10H_2_O, 2 g CaCl_2_·2H_2_O, and 0.1 g (NH_4_)6Mo7O_24_·4H_2_O. When OD_600_ of a culture reached 0.6-0.8 during flask cultivation, 1 mM isopropyl β-D-1-thiogalactopyranoside (IPTG) was added to induce the gene expression. After the IPTG induction, the cells were cultivated for another 48 h.

### Gene cloning and plasmid construction

Standard protocols were used for PCR, gel electrophoresis, and transformation experiments^52^. All the plasmids used in this study are also listed in Supplementary Table 1. Primers used to construct plasmids are listed in Supplementary Table 2. All heterologous genes employed in this study are listed in Supplementary Table 3. DNA sequences of codon-optimized genes are available in Supplementary Table 4. *E. coli* DH10β (New England Biolabs) was used as a host strain for routine gene cloning, in lysogeny broth (LB) medium (per liter: 10 g tryptone, 5 g yeast extract and 10 g NaCl) or on LB agar plates (1.5% agar, w/v) at 37°C supplemented with appropriate antibiotics as described above (Supplementary Table 1). Lamp Pfu DNA polymerase purchased from BIOFACT was used for PCR reactions. Restriction endonucleases and T4 ligase were purchased from Enzynomics or New England Biolabs. Gene knockouts were conducted in *E. coli* BAP1 strain using the previously established protocol^53^.

### SDS-PAGE analysis

To confirm heterologous expression of *cotA* encoding laccase, *E. coli* BAP1Δ*aspC* strain harboring pACN1 was cultured overnight in LB media, and small aliquot was transferred to test tubes containing 10 mL R/2 medium with 20 g/L of glucose. The cells were grown at 30°C, and were induced with 1 mM IPTG when OD_600_ of the culture reached 0.6-0.8. Cells were additionally grown for around 6 h, and a sample was collected, which had OD_600_ of about 10 when resuspended in 0.3 mL of phosphate-buffered saline (PBS) solution (pH 7.4). Cell pellets were washed with 1 mL of cold PBS solution (pH 7.4), centrifuged at 15,000 g for 1 min at 4°C, and resuspended in 0.3 mL of the same PBS buffer. The resuspended cells were lysed by bacterial protein extraction reagent B-PER (Thermo Fisher Scientific). In order to precipitate insoluble protein fractions and partially disrupted cells, the lysed samples were centrifuged at 15,000 g and 4 °C for 10 min at. Total protein extracts and soluble extracts were loaded on SDS-polyacrylamide gel, and analyzed for protein expression.

### Metabolome analysis of the control and engineered strains

Absolute quantification of metabolites was conducted at Human Metabolome Technologies (HMT) by using capillary electrophoresis time-of-flight mass spectrometry measurement performed (Supplementary Data 1). Strains for metabolome analysis were prepared following the guidelines provided by HMT. Absolute quantification was attempted for 353 metabolites, and 160 metabolites were detected and annotated on the basis of HMT’s standard library and its in-house standard data.

### Fed-batch fermentation

Fed-batch fermentations were conducted using a 6.6 L bioreactor (BioFlo 320, Eppendorf) containing 1.7 L R/2 medium (pH 6.8) supplemented with 20 g/L glucose, and appropriate antibiotics. Cells were inoculated from a colony into a 25 ml test tube containing 10 mL LB medium supplemented with appropriate antibiotics. The cells in the test tubes were cultivated overnight in a rotary shaker at 37°C and 200 rpm. Next, four 250 mL baffled flasks, containing 50 mL of R/2 medium supplemented with 20 g/L glucose and appropriate antibiotics, were inoculated with 2 mL aliquot of the seed culture. The cells were cultured at 30°C and 200 rpm until their OD_600_ reached around 4. A total of 200 mL of the cultured cells were transferred into the bioreactor. The culture pH was controlled at 6.8 by automatic feeding of 28% (v/v) ammonia solution, and the temperature was maintained at 30°C. The dissolved oxygen (DO) level was maintained at 40% of air saturation by supplying air at a rate of 2 L/min. The agitation speed was automatically controlled up to 1000 rpm. When OD_600_ of the culture reached between 65 and 80, which was based on the experimental trials, 1 mM IPTG was added to induce the enzyme expression (Figs. 2-5, 7 and 8). L-tryptophan and L-methionine were also added during the induction when required (Figs. 2, 3 and 4). Only L-methionine was added for the strains engineered for de novo production (Figs. 5, 7 and 8). The pH-stat feeding method was employed to supply exhausted nutrients to the culture. The feeding solution contains the following per liter: 700 g glucose, 5 mL TMS, 8 g MgSO_4_·7H_2_O and appropriate antibiotics. When the pH rose higher than 6.83 due to carbon source exhaustion, the feeding solution was automatically added.

### Measurement of dry cell weight

Cell growth was monitored by measuring the absorbance at 600 nm (OD_600_) with Ultrospec 3100 spectrophotometer (Amersham Biosciences), and dry cell weight was measured via gravimetric analysis. The cell culture was washed by mixing it with water (1:1) and centrifuged, and the supernatant was removed. The cleaning process was carried out twice. The clean samples were dried in an oven at 70°C for 24 h, cooled to room temperature, and the resulting biomass was gravimetrically quantified using an analytical balance.

### Extraction and quantification of chemicals

The standards for L-tryptophan, L-methionine, L-kynurenine, 3-hydroxykynurenine, kynurenic acid, and 3-HA were purchased from Merck (Sigma-Aldrich). For L-kynurenine and kynurenic acid, 1 mL cell culture was centrifuged, and its supernatant was analyzed after filtering using 0.22 μm PTFE syringe filter (FUTECS). All the prepared samples were analyzed using high performance liquid chromatography (HPLC; 1100 Series HPLC; Agilent Technologies) connected with MS (LC/MSD VL; Agilent Technologies). LC-MS was operated with: two mobile phase solvents, solvent A (0.1% formic acid in water) and solvent B (acetonitrile); column, Eclipse plus C18 (5 µm, 4.6 x 150 mm); electrospray ionization (ESI) positive ion mode for MS; and wavelength, 254 nm. The eluent was continuously injected into the MS using ESI positive ion mode with: fragmentor, 80V; drying gas flow, 12.0 L/min; drying gas temperature, 350 °C; nebulizer pressure, 30 psig; capillary voltage, 2.5 kV. For analysis, scan mode was used and the scanned mass range was m/z of 100-500.

3-HMA was purchased from An-gene. For extraction of 3-HMA, actinocin and 3-HA, ethyl acetate extract of the cells was prepared by adding equal amount of ethyl acetate to the cell culture. The mixture was placed on a rocking shaker for 30 min. Next, the mixture was centrifuged at 3500 rpm for 20 min to separate the organic phase from the cell pellet. The separated organic extract was carefully decanted, and kept for drying under air blower to remove ethyl acetate. The dried extract was further dissolved in 1 mL methanol, and filtered through 0.22 μm PTFE syringe filter (FUTECS). The prepared samples were subjected to LC-MS for quantification using the same conditions as mentioned above.

Because the standard was not available for actinocin, a MS peak representing the same m/z as actinocin was further purified using fraction collector (G1364C; Agilent Technologies) attached to HPLC (1260 Infinity II; Agilent Technologies) equipped with DAD detectors (G7115A; Agilent Technologies) (Figs. 4b and 5a). Eclipse plus C18 column (4.6 × 150 mm; Agilent Eclipse plus C18) was used. Mobile phase was run at a flow rate of 0.6 mL/min; the mobile phase consists of solvent A (0.1% (v/v) formic acid in distilled water) and solvent B (acetonitrile). The following gradient was applied: 0-3 min, an isocratic condition at 5% solvent B; 3-16 min, a linear gradient of solvent B from 5% to 70%; and 16-25 min, an isocratic condition at 70% solvent B (all in vol%). Samples were monitored at 254 nm. Concentration of actinocin was determined by drying and weighing the collected sample and mapping the area of HPLC or MS peaks to each calibration curve generated using the purified and diluted samples. The purified sample was further subjected to NMR analysis for confirmation.

### NMR analysis for actinocin

NMR spectra of actinocin were acquired with online liquid chromatography (1260 Infinity HPLC System, Agilent Technologies) connected with solid phase extraction (SPE) system (Bruker) and NMR (Cryo-800 MHz FT-NMR Spectrometer, Bruker) installed at Korea Basic Science Institute (KBSI). First, target HPLC peaks were selected by HPLC connected with MS (Impact HD QTOF Mass Spectrometer, Bruker). Two mobile phase solvents were used: solvent A (0.1% formic acid in water) and solvent B (acetonitrile). The selected MS peak was subjected to the NMR system which was trapped by HySphere Resin GP cartridges (inner diameter of 2 mm and particle size of 10-12 μm) using the Bruker Spark Prospekt 2 SPE unit, after post-column addition of water using a Knauer K-100 HPLC pump. The trapped target compound was dried with nitrogen gas for 30 min, and was eluted with CD_3_CN (acetonitrile-d3) for NMR analysis. ^1^H NMR spectra were acquired from a Bruker AVANCE III HD 800 MHz FT-NMR spectrometer using a 5 mm triple-resonance inverse (TCI) CryoProbe with Z-gradient (Bruker) at 298 K, with chemical shifts reported in parts per million relative to CD_3_CN.

## Data availability

The data supporting the findings of this study are available within the article and its Supplementary Information and Supplementary Data. Additional data are available from the corresponding author upon reasonable request.

## Acknowledgments

This work was supported by the Bio & Medical Technology Development Program (NRF- 2018M3A9H3020459), by the Development of next-generation biorefinery platform technologies for leading bio-based chemicals industry project (2022M3J5A1056072), and by the Development of platform technologies of microbial cell factories for the next-generation biorefineries project (2022M3J5A1056117) from National Research Foundation supported by the Korean Ministry of Science and ICT. This work was also carried out with the support of “Cooperative Research Program for Agriculture Science and Technology Development (RS- 2021-RD009210)” from Rural Development Administration, and the Novo Nordisk Foundation (grant NNF16OC0021746).

## Author contributions

H.U.K. conceived the project. K.S., M.R.G. and G.L. performed experiments. All the authors analyzed data, and wrote the manuscript.

## Additional information

Supplementary information is available for this paper.

## References

1. Savitz, J. The kynurenine pathway: a finger in every pie. Mol. Psychiatry 25, 131–147 (2020).

2. Turski, M. P., Turska, M., Zgrajka, W., Kuc, D. & Turski, W. A. Presence of kynurenic acid in food and honeybee products. Amino Acids 36, 75–80 (2009).

3. Milart, P. et al. Kynurenic acid as the neglected ingredient of commercial baby formulas. Sci. Rep. 9, 6108 (2019).

4. Tomaszewska, E. et al. Chronic dietary supplementation with kynurenic acid, a neuroactive metabolite of tryptophan, decreased body weight without negative influence on densitometry and mandibular bone biomechanical endurance in young rats. PLOS ONE 14, e0226205 (2019).

5. Matsuo, M. et al. Antioxidative mechanism and apoptosis induction by 3-hydroxyanthranilic acid, an antioxidant in Indonesian food Tempeh, in the human hepatoma-derived cell line, HuH-7. J. Nutr. Sci. Vitaminol. (Tokyo) 43, 249–259 (1997).

6. Sutphin, G. L. Systemic elevation of 3-hydroxyanthranilic (3HAA) to extend lifespan and delay alzheimer’s pathology. *Innov*. Aging 2, 74–74 (2018).

7. Krause, D. et al. The tryptophan metabolite 3-hydroxyanthranilic acid plays anti-inflammatory and neuroprotective roles during inflammation: role of hemeoxygenase-1. Am. J. Pathol. 179, 1360–1372 (2011).

8. Torrens-Spence, M. P., Liu, C.-T. & Weng, J.-K. Engineering new branches of the kynurenine pathway to produce oxo-(2-aminophenyl) and quinoline scaffolds in yeast. ACS Synth. Biol. 8, 2735–2745 (2019).

9. Robbel, L. & Marahiel, M. A. Daptomycin, a bacterial lipopeptide synthesized by a nonribosomal machinery. J. Biol. Chem. 285, 27501–27508 (2010).

10. Liao, G. et al. Manipulation of kynurenine pathway for enhanced daptomycin production in *Streptomyces roseosporus*. Biotechnol. Prog. 29, 847–852 (2013).

11. Lyu, Z.-Y. et al. Improving the yield and quality of daptomycin in *Streptomyces roseosporus* by multilevel metabolic engineering. Front. Microbiol. 13, 872397 (2022).

12. Yamanaka, K. et al. Direct cloning and refactoring of a silent lipopeptide biosynthetic gene cluster yields the antibiotic taromycin A. Proc. Natl. Acad. Sci. 111, 1957–1962 (2014).

13. Praseuth, A. P. et al. Complete sequence of biosynthetic gene cluster responsible for producing triostin A and evaluation of quinomycin-type antibiotics from *Streptomyces triostinicus*. Biotechnol. Prog. 24, 1226–1231 (2008).

14. Sato, M., Nakazawa, T., Tsunematsu, Y., Hotta, K. & Watanabe, K. Echinomycin biosynthesis. Curr. Opin. Chem. Biol. 17, 537–545 (2013).

15. Lombó, F. et al. Deciphering the biosynthesis pathway of the antitumor thiocoraline from a marine actinomycete and its expression in two streptomyces species. *Chembiochem Eur*. J. Chem. Biol. 7, 366–376 (2006).

16. Chen, F.-M. The nature of actinomycin D binding to d(AACCAXYG) sequence motifs. Nucleic Acids Res. 32, 271–277 (2004).

17. Golub, E. E., Ward, M. A. & Nishimura, J. S. Biosynthesis of the actinomycin chromophore: Incorporation of 3-hydroxy-4-methylanthranilic acid into actinomycins by *Streptomyces antibioticus*. J. Bacteriol. 100, 977–984 (1969).

18. Troost, T. & Katz, E. Phenoxazinone biosynthesis: accumulation of a precursor, 4-methyl-3-hydroxyanthranilic acid, by mutants of *Streptomyces parvulus*. J. Gen. Microbiol. 111, 121–132 (1979).

19. Li, W., Khullar, A., Chou, S., Sacramo, A. & Gerratana, B. Biosynthesis of sibiromycin, a potent antitumor antibiotic. Appl. Environ. Microbiol. 75, 2869–2878 (2009).

20. Giessen, T. W., Kraas, F. I. & Marahiel, M. A. A four-enzyme pathway for 3,5-dihydroxy-4-methylanthranilic acid formation and incorporation into the antitumor antibiotic sibiromycin. Biochemistry 50, 5680–5692 (2011).

21. Hu, Y., Phelan, V. V., Farnet, C. M., Zazopoulos, E. & Bachmann, B. O. Reassembly of anthramycin biosynthetic gene cluster by using recombinogenic cassettes. *Chembiochem Eur*. J. Chem. Biol. 9, 1603–1608 (2008).

22. Umetsu, S., Kanda, M., Imai, I., Sakai, R. & Fujita, M. J. Questiomycins, algicidal compounds produced by the marine bacterium *Alteromonas* sp. D and their production cue. Mol. Basel Switz. 24, 4522 (2019).

23. Ezeokonkwo, M. A. et al. Synthesis and Antimicrobial Activity of New Derivatives of Angular Polycyclic Phenoxazine Ring System. Orient. J. Chem. 35, 1320–1326 (2019).

24. Eggert, C., Temp, U., Dean, J. F. D. & Eriksson, K.-E. L. Laccase-mediated formation of the phenoxazinone derivative, cinnabarinic acid. FEBS Lett. 376, 202–206 (1995).

25. Yue, S.-J. et al. Synthesis of cinnabarinic acid by metabolically engineered *Pseudomonas chlororaphis* GP72. Biotechnol. Bioeng. 116, 3072–3083 (2019).

26. Keller, U., Lang, M., Crnovcic, I., Pfennig, F. & Schauwecker, F. The actinomycin biosynthetic gene cluster of *Streptomyces chrysomallus*: a genetic hall of mirrors for synthesis of a molecule with mirror symmetry. J. Bacteriol. 192, 2583–2595 (2010).

27. Crnovčić, I. et al. Genetic interrelations in the actinomycin biosynthetic gene clusters of *Streptomyces antibioticus* IMRU 3720 and Streptomyces chrysomallus ATCC11523, producers of actinomycin X and actinomycin C. Adv. Appl. Bioinforma. Chem. AABC 10, 29–46 (2017).

28. Ohashi, K., Kawai, S. & Murata, K. Secretion of quinolinic acid, an intermediate in the kynurenine pathway, for utilization in NAD+ biosynthesis in the yeast *Saccharomyces cerevisiae*. Eukaryot. Cell 12, 648–653 (2013).

29. Alberati-Giani, D. et al. Cloning and functional expression of human kynurenine 3-monooxygenase. FEBS Lett. 410, 407–412 (1997).

30. Walther, T. et al. Construction of a synthetic metabolic pathway for the production of 2,4- dihydroxybutyric acid from homoserine. Metab. Eng. 45, 237–245 (2018).

31. Han, Q., Fang, J. & Li, J. Kynurenine aminotransferase and glutamine transaminase K of *Escherichia coli*: identity with aspartate aminotransferase. Biochem. J. 360, 617–623 (2001).

32. Kuramitsu, S., Okuno, S., Ogawa, T., Ogawa, H. & Kagamiyama, H. Aspartate aminotransferase of *Escherichia coli*: nucleotide sequence of the aspC gene. J. Biochem. (Tokyo) 97, 1259–1262 (1985).

33. Fontecave, M., Atta, M. & Mulliez, E. S-adenosylmethionine: nothing goes to waste. Trends Biochem. Sci. 29, 243–249 (2004).

34. Markham, G. D., DeParasis, J. & Gatmaitan, J. The sequence of metK, the structural gene for S-adenosylmethionine synthetase in *Escherichia coli*. J. Biol. Chem. 259, 14505– 14507 (1984).

35. Moccand, C., Kaufmann, M. & Fitzpatrick, T. B. It takes two to tango: Defining an essential second active Site in pyridoxal 5′-phosphate synthase. PLoS ONE 6, e16042 (2011).

36. Mittenhuber, G. Phylogenetic analyses and comparative genomics of vitamin B6 (pyridoxine) and pyridoxal phosphate biosynthesis pathways. J. Mol. Microbiol. Biotechnol. 3, 1–20 (2001).

37. Ma, W. et al. Engineering a pyridoxal 5’-phosphate supply for cadaverine production by using *Escherichia coli* whole-cell biocatalysis. Sci. Rep. 5, 15630 (2015).

38. Yang, J. G. & Rees, D. C. The allosteric regulatory mechanism of the *Escherichia coli* MetNI Methionine ATP Binding Cassette (ABC) Transporter. J. Biol. Chem. 290, 9135–9140 (2015).

39. Ravi Kant, H., Balamurali, M. & Meenakshisundaram, S. Enhancing precursors availability in *Pichia pastoris* for the overproduction of S-adenosyl-L-methionine employing molecular strategies with process tuning. J. Biotechnol. 188, 112–121 (2014).

40. Wang, C. et al. Cloning and characterization of CotA laccase from *Bacillus subtilis* WD23 decoloring dyes. Ann. Microbiol. 66, 461–467 (2016).

41. Osiadacz, J., Al-Adhami, A. J. H., Bajraszewska, D., Fischer, P. & Peczyñska-Czoch, W. On the use of *Trametes versicolor* laccase for the conversion of 4-methyl-3- hydroxyanthranilic acid to actinocin chromophore. J. Biotechnol. 72, 141–149 (1999).

42. Du, J., Yang, D., Luo, Z. W. & Lee, S. Y. Metabolic engineering of *Escherichia coli* for the production of indirubin from glucose. J. Biotechnol. 267, 19–28 (2018).

43. Ikeda, M. Towards bacterial strains overproducing l-tryptophan and other aromatics by metabolic engineering. Appl. Microbiol. Biotechnol. 69, 615–626 (2006).

44. Zhao, Z.-J. et al. Development of l-tryptophan production strains by defined genetic modification in *Escherichia coli*. J. Ind. Microbiol. Biotechnol. 38, 1921–1929 (2011).

45. Rodriguez, A., Kildegaard, K. R., Li, M., Borodina, I. & Nielsen, J. Establishment of a yeast platform strain for production of p-coumaric acid through metabolic engineering of aromatic amino acid biosynthesis. Metab. Eng. 31, 181–188 (2015).

46. Li, G. & Young, K. D. Indole production by the tryptophanase TnaA in *Escherichia coli* is determined by the amount of exogenous tryptophan. Microbiol. Read. Engl. 159, 402– 410 (2013).

47. Liu, J. & Summers, D. Indole at low concentration helps exponentially growing *Escherichia coli* survive at high temperature. PLOS ONE 12, e0188853 (2017).

48. Rakicka-Pustułka, M. et al. The microbial production of kynurenic acid using *Yarrowia lipolytica* yeast growing on crude glycerol and soybean molasses. Front. Bioeng. Biotechnol. 10, 936137 (2022).

49. Yue, S. et al. Synthesis of cinnabarinic acid by metabolically engineered *Pseudomonas chlororaphis* GP72. Biotechnol. Bioeng. 116, 3072–3083 (2019).

50. Niu, H. et al. Metabolic engineering for improving L-tryptophan production in *Escherichia coli*. J. Ind. Microbiol. Biotechnol. 46, 55–65 (2019).

51. Pfeifer, B., Hu, Z., Licari, P. & Khosla, C. Process and metabolic strategies for improved production of *Escherichia coli*-derived 6-deoxyerythronolide B. Appl. Environ. Microbiol. 68, 3287–3292 (2002).

52. Green, R. & Rogers, E. J. Transformation of chemically competent E. coli. in Methods in Enzymology vol. 529 329–336 (Elsevier, 2013).

53. Song, C. W. & Lee, S. Y. Rapid one-step inactivation of single or multiple genes in *Escherichia coli*. Biotechnol. J. 8, 776–784 (2013).

