## Supplementary Information for "De novo production of an antitumor precursor actinocin and other medicinal molecules from kynurenine pathway in *Escherichia coli*"

### Supplementary Tables

**Supplementary Table 1 | List of strains and plasmids used in this study.**

| Strain/Plasmid | Description | Source |
|---|---|---|
| **Strains** | | |
| *Escherichia coli* DH10β | *Δ*(*ara-leu*) *7697 araD139 fhuA ΔlacX74 galK16 galE15* e14- ϕ*80dlacZΔM15recA1relA1 endA1 nupGrpsL* (StrR) *rph spoT1Δ*(*mrr-hsdRMS-mcrBC*) | New England Biolabs |
| *E. coli* BAP1 | BL21(DE3) ΔprpRBCD::PT7-sfp-PT7-prpE | 1 |
| *E. coli* BAP1Δ*aspC* | BL21(DE3) ΔprpRBCD::PT7-sfp-PT7-prpE Δ*aspC* | This study |
| *E. coli* BAP1Δ*aspC*Δ*tnaA* | BL21(DE3) ΔprpRBCD::PT7-sfp-PT7-prpE Δ*aspC* Δ*tnaA* | This study |
| **Plasmids** | | |
| pCDFDuet-1 | $Spc^R$, T7 promoter, CDF origin | Novagen |
| pRSFDuet-A | $Ap^R$, T7 promoter, RSF origin | Novagen |
| pET-30a | $Km^R$, T7 promoter, pBR322 origin | Novagen |
| pTac15K | $Km^R$, Tac promoter, p15A origin | Novagen |
| pACYCDtac | $Cm^R$, Tac promoter, p15A origin | Novagen |
| pACYCDtac-rsf | $Cm^R$, Tac promoter, RSF origin | This study |
| pECmulox | $Ap^R$, loxLE- $Cm^R$ gene-loxRE | 2 |
| pCW611 | $Ap^R$, λ-Red recombinase under arabinose-inducible PBAD promoter, Cre-recombinase under IPTG inducible lacUV5 promoter, ts origin | 2 |
| pG | pCDFDuet-1 derivative containing *acmG* from *Streptomyces anulatus* | This study |
| pF | pCDFDuet-1 derivative containing *acmF* from *S. anulatus* | This study |

| | | |
|---|---|---|
| psaK | pCDFDuet-1 derivative containing *KMO* from *S. anulatus* | This study |
| pC | pCDFDuet-1 derivative containing codon-optimized *sibC* from *Streptosporangium sibiricum* | This study |
| pB | pCDFDuet-1 derivative containing codon-optimized *bna4* from *Saccharomyces cerevisiae* | This study |
| phsK | pCDFDuet-1 derivative containing codon-optimized *hsKMO* from *Homo sapiens* | This study |
| pI | pCDFDuet-1 derivative containing *acmI* from *S. anulatus* | This study |
| paL | pCDFDuet-1 derivative containing *acmL* from *S. anulatus* | This study |
| pL | pCDFDuet-1 derivative containing codon-optimized *sibC S. sibiricum* | This study |
| pH | pCDFDuet-1 derivative containing *acmH* from *S. anulatus* | This study |
| pH` | pCDFDuet-1 derivative containing $NH_6$*acmH* (encoding N-terminus His6-tagged AcmH) from *S. anulatus* | This study |
| pK` | pCDFDuet-1 derivative containing $NH_6$*acmK* (encoding N-terminus His6-tagged AcmK) from *S. anulatus* | This study |
| pQ | pCDFDuet-1 derivative containing codon-optimized *sibQ* from *S. sibiricum* | This study |
| pKN1 | pCDFDuet-1 derivative containing *acmG* and *acmF* from *S. anulatus* | This study |
| pKN2 | pCDFDuet-1 derivative containing *acmG* and $NH_6$*acmF* (encoding N-terminus His6-tagged AcmF) from *S. anulatus* | This study |
| pKN3 | pCDFDuet-1 derivative containing $NH_6$*acmG* (encoding N-terminus His6-tagged AcmG) and $NH_6$*acmF* (encoding N-terminus His6-tagged AcmF) from *S. anulatus* | This study |
| pGF`saK | pCDFDuet-1 derivative containing *acmG* and $NH_6$*acmF* (encoding N-terminus His6-tagged AcmF) and *saKMO* from *S. anulatus* | This study |
| psaK`trxA | pCDFDuet-1 derivative containing *saKMO`trxA* (encoding N-terminus His6-tagged saKMO and TrxA protein attached at C-terminal) | This study |
| pHK1 | pCDFDuet-1 derivative containing *acmG* and $NH_6$*acmF* (encoding N-terminus His6-tagged AcmF) and *saKMO`trxA* (encoding N-terminus His6-tagged saKMO and TrxA protein attached at C-terminal) | This study |
| pHK2 | pCDFDuet-1 derivative containing *acmG* and $NH_6$*acmF* (encoding N-terminus His6- | This study |

| | | |
|---|---|---|
| | tagged AcmF) from *S. anulatus* and codon-optimized *sibC S. sibiricum* | |
| pHK3 | pCDFDuet-1 derivative containing *acmG*, $NH_6$*acmF* (encoding N-terminus His6-tagged AcmF) from *S. anulatus* and codon-optimized *bna4 S. cerevisiae* | This study |
| pHK4 | pCDFDuet-1 derivative containing *acmG*, $NH_6$*acmF* (encoding N-terminus His6-tagged AcmF) from *S. anulatus* and codon-optimized *hsKMO* from *H. sapiens* | This study |
| pHMA1 | pCDFDuet-1 derivative containing *acmG*, $NH_6$*acmF* (encoding N-terminus His6-tagged AcmF), *acmI* and $NH_6$*acmH* (encoding N-terminus His6-tagged AcmH) from *S. anulatus*; codon-optimized *sibC* from *S. sibiricum* | This study |
| pHMA2 | pCDFDuet-1 derivative containing *acmG*, $NH_6$*acmF* (encoding N-terminus His6-tagged acmF), *acmI* and $NH_6$*acmH* (encoding N-terminus His6-tagged AcmH) from *S. anulatus*; codon-optimized *bna4* from *S. cerevisiae* | This study |
| pHMA3 | pCDFDuet-1 derivative containing *acmG*, $NH_6$*acmF* (encoding acmF attached with N-terminal 6×His-tag), *acmL* and $NH_6$*acmH* (encoding N-terminus His6-tagged AcmH) from *S. anulatus*; codon-optimized *sibC* from *S. sibiricum* | This study |
| pHMA4 | pCDFDuet-1 derivative containing *acmG*, $NH_6$*acmF* (encoding N-terminus His6-tagged AcmF), *acmL* and $NH_6$*acmH* (encoding N-terminus His6-tagged AcmH) from *S. anulatus*; codon-optimized *bna4* from *S. cerevisiae* | This study |
| pHMA5 | pCDFDuet-1 derivative containing *acmG*, $NH_6$*acmF* (encoding N-terminus His6-tagged AcmF), and $NH_6$*acmH* (encoding N-terminus His6-tagged AcmH) from *S. anulatus*; codon-optimized *sibC* and *sibL* from *S. sibiricum* | This study |
| pHMA6 | pCDFDuet-1 derivative containing *acmG*, $NH_6$*acmF* (encoding N-terminus His6-tagged AcmF), and $NH_6$*acmH* (encoding N-terminus His6-tagged AcmH) from *S. anulatus*; codon-optimized *bna4* from *S. cerevisiae;* codon-optimized *sibL* from *S. sibiricum* | This study |
| pHMA7 | pCDFDuet-1 derivative containing *acmG*, $NH_6$*acmF* (encoding N-terminus His6-tagged AcmF, and $NH_6$*acmK* (encoding N-terminus His6-tagged AcmK) from *S. anulatus*; codon-optimized *sibC* and *sibL* from *S. sibiricum* | This study |
| pHMA8 | pCDFDuet-1 derivative containing *acmG*, $NH_6$*acmF* (encoding N-terminus His6-tagged AcmF), and $NH_6$*acmK* (encoding N-terminus His6-tagged AcmK) from *S. anulatus*; codon-optimized *bna4* from *S. cerevisiae;* codon-optimized *sibL* from *S. sibiricum* | This study |
| pHMA9 | pCDFDuet-1 derivative containing *acmG*, $NH_6$*acmF* (encoding N-terminus His6-tagged AcmF) from *S. anulatus*; codon-optimized *sibC, sibL* and *sibQ* from *S. sibiricum* | This study |

| | | |
|---|---|---|
| pHMA10 | pCDFDuet-1 derivative containing *acmG*, $NH_6$*acmF* (encoding N-terminus His6-tagged AcmF) from *S. anulatus*; codon-optimized *bna4* from *S. cerevisiae,* and *sibL* and *sibQ* from *S. sibiricum* | This study |
| pSAM | pET-30a derivative containing *metK* from *E. coli* | This study |
| pSP1 | pET-30a derivative containing *metK* and *pdxJ* from *E. coli* | This study |
| pSP2 | pET-30a derivative containing *metK*, *pdxJ* and *dxs* from *E. coli* | This study |
| pSAM1 | pET-30a derivative containing *metK* and *metNIQ* from *E. coli* | This study |
| pSAM2 | pET-30a derivative containing *metK* from *E. coli* and codon-optimized *mup1* from *S. cerevisiae* | This study |
| pSAM3 | pET-30a derivative containing *metK* and *adk* from *E. coli* and codon-optimized *mup1* from *S. cerevisiae* | This study |
| pSAM4 | Modified pET-30a derivative containing *metK* and *adk* from *E. coli* and codon-optimized *mup1* from *S. cerevisiae*, and $Km^R$ marker replaced with $Ap^R$ resistance marker | This study |
| pJ | pRSFDuet-A derivative containing *pdxJ* from *E. coli* | This study |
| pPLP | pRSFDuet-A derivative containing *pdxJ* and *dxs* from *E. coli* | This study |
| pACN1 | pRSFDuet-A derivative containing *cotA* from *Bacillus subtilis* 168 | This study |
| pACN2 | Modified pET-30a derivative containing *metK* and *adk* from *E. coli*, codon-optimized *mup1* from *S. cerevisiae* and *cotA* from *B. subtilis* 168 and Km marker replaced with $Ap^R$ resistance marker | This study |
| pTRP | pTac15K derivative containing *aroG*$^{fbr}$ (encoding $AroG^{A146N}$), *trpE*$^{fbr}$ (encoding $TrpE^{S40F}$) and *aroL* from *E. coli* | 3 |
| pKYNA | pCDFDuet-1 derivative containing *acmG* and $NH_6$*acmF* (encoding N-terminus His6-tagged AcmF) from *S. anulatus* and *aspC* from *E. coli* | This study |
| pHA1 | pCDFDuet-1 derivative containing *acmG*, $NH_6$*acmF* (encoding N-terminus His6-tagged AcmF) from *S. anulatus*; codon-optimized *sibC* and *sibQ* from *S. sibiricum* | This study |
| pTRP1 | Rearranged pTac15K derivative containing *aroG*$^{fbr}$ (encoding $AroG^{A146N}$), *aroL* and *trpE*$^{fbr}$ (encoding $TrpE^{S40F}$) from *E. coli* | This study |
| pFbaA | pACYCDtac-rsf derivative containing *fbaA* form *E. coli* | This study |

| | | |
|---|---|---|
| pIcd | pACYCDtac-rsf derivative containing *icd* from *E. coli* | This study |
| pIF | pACYCDtac-rsf derivative containing *fbaA* and *icd* from *E. coli* | This study |

Abbreviations: $Ap^R$, ampicillin resistance; $Km^R$, kanamycin resistance; $Cm^R$, chloramphenicol resistance; $Spc^R$, spectinomycin resistance; $^{fbr}$, feedback inhibition eliminated by point mutation.

**Supplementary Table 2 | List of primers used in the study.**

| Primer name | Sequence (5`→3`) | Purpose |
|---|---|---|
| acmG_gib_F1 | gtttaactttaataaggagatatatgaccgtcgaacaggaagccac | pG construction (Gibson assembly) |
| acmG_gib_R1 | ggtgatggctgctgcccatggttcagatctccgagcgcaccgacc | |
| pCDF_BB_R1 | atatctccttattaaagttaaacaaaattatttctacagggga | |
| pCDF_BB_F1 | gtgtggcgctttctcatagctca | |
| pCDF_BB_R2 | ccgagccgagataccagtgtgtgagc | |
| pCDF_BB_F2 | accatgggcagcagccatcac | |
| acmF_gib_F1 | gtttaactttaataaggagatatatgaccggcgccgccgtgtaccg | pF construction (Gibson along with pCDF backbone, above primers) |
| acmF_gib_R1 | ggtgatggctgctgcccatggttcatgcggtcgcctcctccggttc | |
| acmG_EcoR1_F | gcttcaattcatctctGAATTCatgaccgtcgaacaggaagccac | pGF construction (Restriction digestion and ligation with pCDF backbone) |
| acmG_RBS_R | atgtatatctccttcttatacttaactatcagatctccgagcgcaccgacc | |
| acmF_RBS_F | tagttaagtataagaaggagatatacatatgaccggcgccgccgtgtaccg | |
| acmF_Sac1_R | gcatcatactacGAGCTCtcatgcggtcgcctcctccggttc | |
| acmG_mod_F1 | accgtcgaacaggaagccac | pG` construction (Insertion of 6xHIS tag in pG plasmid by Gibson assembly) |
| pCDF_BB_R2 | ccgagccgagataccagtgtgtgagc | |
| pCDF_hisG_R1 | acaccccgtggcttcctgttcgacggtatggtgatggtgatggtgcatatatctccttattaaagttaaacaaaattatttc | |
| pCDF_BB_F1 | gtgtggcgctttctcatagctca | |
| acmF_mod_F1 | accggcgccgccgtgtaccg | pF` construction (Insertion of 6xHIS tag in pF plasmid by Gibson assembly) |
| pCDF_hisF_R2 | tgccgcggtacacggcggcgccggtatggtgatggtgatggtgcatatatctccttattaaagttaaacaaaattatttc | |
| pCDF_hisF_R1 | tgccgcggtacacggcggcgccggtatggtgatggtgatggtgcatatgtatatctccttcttatacttaactatcag | pG`F` construction (Gibson assembly) |
| sibC_pCDF_F1 | gtatattagttaagtataagaaggagatatacatatgtccgcgacaccacctaagg | pGF`C construction (Gibson assembly) |
| sibC_pCDF_R1 | ggccgatatccaattgagatctgccattcaatgctccccctttgcaggg | |

| | | |
|---|---|---|
| gib_pCDF_R2 | atgtatatctccttcttatacttaactaatatactaagatgggg | |
| gib_pCDF_F2 | atggcagatctcaattggatatcgg | |
| bna4_pCDF_F1 | gtatattagttaagtataagaaggagatatacatatgagtgaatctgtcgcgatcatcgg | pGF`B construction (Gibson assembly) |
| bna4_pCDF_R1 | ggccgatatccaattgagatctgccattcatgagcgctcccgcgtcaa | |
| KMO_mod_F2 | gtagaaataattttgtttaactttaataaggagatatatgcatcatcatcatcatcataacgtcgtcgtcgtcggggcg | psaK`trxA construction (Gibson assembly) |
| KMO_trxA_R1 | cagtcaggtgaataattttatcgctcatgcatcttccgctccgcttcc | |
| KMO_trxA_F1 | gagttcctcgacgctaacctggcgtaaaccatgggcagcagccatcac | |
| trxA_R1 | ttacgccaggttagcgtcgaggaac | |
| trxA_F1 | atgagcgataaaattattcacctgactgacg | |
| H-KMO_pCDF_F1 | gtatattagttaagtataagaaggagatatacatatggactcttccgtcatccagcg | pGF`hsK construction (Gibson assembly) |
| H-KMO_pCDF_R1 | ggccgatatccaattgagatctgccattcaacgactaatcaaattagaaatctgttccaggc | |
| sibC_mod_F2 | gtatattagttaagtataagaaggagatatacatatgcaccatcaccatcaccatcaccattccgcgacaccacctaaggccgt | pGF`C` construction (Gibson assembly) |
| pCDF_hisSib_R1 | gattacggccttaggtggtgtcgcggaatggtgatggtgatggtgcatatgtatatctccttcttatacttaactaa | |
| sibC_RBS_R1 | atgtatatctccttcttaaagttaaacatcaatgctcccccttttgcag | pGF`CIH` construction (Gibson assembly) |
| acmI_RBS_F1 | tgatgtttaactttaagaaggagatatacatatggcagacgtccgccccttcg | |
| acmI_RBS_R1 | atgtatatctccttcttatacttaactatcacttgaccgcgctgatcacgc | |
| acmH_mod_F2 | gtgatagttaagtataagaaggagatatacatatgcaccatcaccatcaccatcaccataacggcgggacaacccgtgag | |
| acmH_gib_R1 | cgatatccaattgagatctgccattcaggtcacccgtgccagttcg | |
| bna4_RBS_R1 | atgtatatctccttcttaaagttaaacatcatgagcgctcccgcgtcaa | pGF`BIH` construction (Gibson assembly) |
| bna4_pCDF_F1 | gtatattagttaagtataagaaggagatatacatatgagtgaatctgtcgcgatcatcgg | |
| acmL_RBS_F1 | tgatgtttaactttaagaaggagatatacatatgcctcacgcgtccccgctca | pGF`CaLH` and pGF`BaLH` construction (Gibson assembly) |
| acmL_RBS_R1 | catatgtatatctccttcttatacttaactatcacttgaccgcgctgatcacgcc | |
| sibL_RBS_F1 | tgtttaactttaagaaggagatatacatatgtcgagcgaattatcggctttag | pGF`CLH` and pGF`BLH` construction (Gibson assembly) |
| sibL_RBS2_R1 | atgtatatctccttcttatacttaactatcaccgtggacgcgagccgctcac | |

| | | |
|---|---|---|
| his-acmK_RBS_F1 | tgatgtttaactttaagaaggagatatacatatgcaccatcaccatcaccatgacgccgaggatttccccaccaac | pGF`CLK` and pGF`BLK` construction (Gibson assembly) |
| acmK_pCDF_R1 | cggccgatatccaattgagatctgccattcaggtcacccgggcgaggcc | |
| sibQ_RBS_F1 | gttgtgagcggctcgcgtccacggtgatgtttaactttaagaaggagatatacatatggaatctacacgtggctgggcag | pGF`CLQ and pGF`BLQ construction (Gibson assembly) |
| sibQ_pCDF_R1 | ccgatatccaattgagatctgccattcaacgcagaacctcccgaag | |
| pdxJ_pRSF_F1 | gtagaaataattttgtttaactttaataaggagatatatgacacttgaaaaatttgtggatgctc | pJ construction (Gibson assembly) |
| pdxJ_pRSF_R1 | gatgatggtgatggctgctgcccatggtttatttatggggatcagttatatcc | |
| pCDF_BB_R1 | atatctccttattaaagttaaacaaaattatttctacagggga | |
| gib_pRSF_F1 | gcagccattggtaactgatttagagg | |
| gib_pRSF_R1 | gtgcataacttcaagacaaagtcctc | |
| pCDF_BB_F2 | accatgggcagcagccatcac | |
| H50-aspC-ko_F1 | gccagtaaaacgtactactcgcgcttatcgtgaagaagcaattattaaatgtatcatacacatacgatttaggtg | Knock-out of *aspC* |
| H50-aspC-ko_R1 | aggtaaaggttaatcctttaatattcgccgcgttctctttcacatgtttacactatagggagaccggcagatc | |
| H100-aspC-ko_F1 | atggaaaactttaaacatctccctgaaccgttccgcattcgtgttattgagccagtaaaacgtactactcgcgct | |
| H100-aspC-ko_R1 | ttaaacttctttcagttttgcggtgaagtgacgcaatacttttggttcgtaggtaaaggttaatcctttaatatt | |
| tnaA-ko-H50-F2 | aatattcacagggatcactgtaattaaaataaatgaaggattatgtaatgtaggtgacactatagaacgcg | Knock-out of *tnaA* |
| tnaA-ko-H50-R2 | acatccttatagccactctgtagtattaattaaacttctttcagttttgctagtggatctgatgggtacc | |
| tnaA-ko-H100-F2 | atgatggtgcttgtatatatacatggcgaattaatggatattgcagatgtaatattcacagggatcactgtaatt | |
| tnaA-ko-H100-R2 | tagaggaaggctatttttgttattgaggatgtagggtaagagagtggctaacatccttatagccactctgtagta | |
| metK_pET_F1 | gtttaactttaagaaggagatatacatatggcaaaacacctttttacgtcc | pSAM construction (Gibson assembly) |
| metK_pET_R1 | cagaagaatgatgatgatgatggtgcatttacttcagaccggcagcatcg | |
| gib_pET_F1 | atgcaccatcatcatcatcattcttctg | |
| BB_pET_R | cattaacgcttctggagaaactcaacgag | |
| BB_pET_F | ctcgttgagtttctccagaagcgttaatg | |
| gib_pET_R1 | atgtatatctccttcttaaagttaaacaaaattatttctagagg | |
| metK_RBS_R1 | atgtatatctccttcttaaagttaaacattacttcagaccggcagcatcg | pSAM1 construction (Gibson |

| | | |
|---|---|---|
| metN_RBS_F1 | tgtttaactttaagaaggagatatacatatgataaaactttcgaatatcacc | assembly) |
| metQ_pET_R1 | gaccagaagaatgatgatgatgatggtgcatttaccagcctttaacagcgccg | |
| mup1_F1 | atgtcagaggggcggaccttcc | pSAM2 construction (Gibson assembly) |
| mup1_R1 | ttacagtgatttctcttgttcagacttgtag | |
| mup1_RBS_F1 | tgtttaactttaagaaggagatatacatatgtcagaggggcggaccttcc | |
| mup1_pET_R1 | ccagaagaatgatgatgatgatggtgcatttacagtgatttctcttgttcagac | |
| adk_F1 | atgcgtatcattctgcttggcgc | pSAM3 construction (Gibson assembly) |
| adk_R1 | ttagccgaggatttttccagatcagc | |
| adk_RBS2_F1 | tagttaagtataagaaggagatatacatatgcgtatcattctgcttggcgc | |
| adk_pET_R1 | cagaagaatgatgatgatgatggtgcatttagccgaggatttttccagatc | |
| cotA_pRSF_F1 | gtagaaataattttgtttaactttaataaggagatatatgacacttgaaaaatttgtggatgctc | pACN1 construction (Gibson assembly) |
| cotA_pRSF_R1 | gatgatggtgatggctgctgcccatggtttatttatggggatcagttatatcc | |
| pCDF_BB_R1 | atatctccttattaaagttaaacaaaattatttctacagggga | |
| gib_pRSF_F1 | gcagccattggtaactgatttagagg | |
| gib_pRSF_R1 | gtgcataacttcaagacaaagtcctc | |
| pCDF_BB_F2 | accatgggcagcagccatcac | |

All the restriction digestion sites are written in capital letters.

N-terminus $His_6$ tags are underlined.

**Supplementary Table 3 | List of heterologous enzymes and their corresponding genes used in this study.**

| Enzyme | Gene | Source organism |
|---|---|---|
| Tryptophan 2,3-dioxygenase | *acmG* | *Streptomyces anulatus* |
| Kynurenine formamidase | *acmF* | *Streptomyces anulatus* |
| Kynurenine 3-monooxygenase | *saKMO* | *Streptomyces anulatus* |
| | *sibC* | *Streptosporangium sibiricum* |
| | *bna4* | *Saccharomyces cerevisiae* |
| | *hsKMO* | *Homo sapiens* |
| Methyltransferase | *acmI* | *Streptomyces anulatus* |
| | *acmL* | *Streptomyces anulatus* |
| | *sibL* | *Streptosporangium sibiricum* |
| Kynureninase | *acmH* | *Streptomyces anulatus* |
| | *acmK* | *Streptomyces anulatus* |
| | *sibQ* | *Streptosporangium sibiricum* |
| Laccase | *cotA* | *Bacillus subtilis* 168 |
| Methionine permease | *mup1* | *Saccharomyces cerevisiae* |

**Supplementary Table 4 | DNA sequence of codon-optimized genes in this study.**

| |
|---|
| **Codon-optimized *sibC* from *Streptosporangium sibiricum*** |
| atgtccgcgacaccacctaaggccgtaatcgtaggggcggggccagtcggttgcttattagcggtcgagttacgccgtcgtgacttcgatgtcgaaatctatgagaaatgcgacgccgatac<br>agtattacgccctggggtcggccggtcattcaaccttacactgacgcaccggggtctgagcgtgctccacccgtggctgcgggaccgcgtttacgaaatcggttcggtgcttcgtcagcgga<br>tcgtacaccataccgatgggaccttgacgcgtcagccttatgggatttccgacgaccaacacctgctctctgtgccacgtcgtgagttgcagcgcattctgttagcggaggcacgcgctagtg<br>gggcacggatgttcttcggtcacagctgcgtaggggcagatgcccatcgcggggaggcgatgtttgttgatccacggggcactgtccgccatgctgcaggtgacatccttatcggttgtgac<br>ggggctaacagcgcgcttcggtacgaactctccaaagggggcgcacgtatgcaggtacatcaacggtatattgcgcatggccacgtagaattgacgctcaccccagaaggcagcgccgct<br>ttagcaccagatggcatgcacttgtggccgcgctctgaccatttccttcaagcccagcctaaccgcgatgggagtacgactactactttgtttatgcctgtagggaatgaaggtggcgagttgc<br>ggtttcggtcactcactggtacggaggcagtccgcgagcattttgaacgtgagtaccctgatatcgcaccacaccttcctttagcagcagatgaagttattcgggctcgcccggctctcttgaaa<br>gtggtagactgcgccccatacaactacgggcgcgctgtgttggtcggcgactccgcacacactattgttccgttctttgggcagggcattaattgctctttcgaggacgcagacatcttatgtcg<br>gctcttagataaatattggcgcagcggcgtagatcgccacgaggccatccgtgcggctggtgcagaattctcagaacagcgtgtagccgccggccaggccattagtcagctttcgatggata<br>atctgcatgaactgtcgaaatccatcgatgatgccgggtttcatcgccggcagcgcttggagcgcatgctccacgaataccgtccagccgagtttacgcaactgtaccagttggtagcgttca<br>cacgcacgccgtacgatgaaattattcgccgtagccgtattgacaaattagcactggatacattgtgcgaactttatgatccggataccgaagcagaaacgattatttcgagttacgcagaggtc<br>cgtcgcggcctggacgaacatgatacgcgggagcctgcgttagcactgacggcaacccctgcaaagggggagcattga |
| **Codon-optimized *bna4* from *Saccharomyces cerevisiae*** |
| atgagtgaatctgtcgcgatcatcggcgccgggctggtaggctgcttggcggcactcgcattttctaaggagggttacaacgtgacattgtacgactttcgccaggatccgcgcttggacaca |

acgaaaaacaagaacctcaagtcaattaatcttgcaatctcggctcgggggattgacgcgttgaagtcaatcgatcctgacgcatgcgaacacattctgcaagacatgatccctatgaaaggc
cgtatgatccacgacctcaagggtcgtcaggaaagtcagttatatggtctgcacggggaggcgatcaacagtatcaaccggtctgtactgaacaattcactgcttgatgaactggaaaaaagc
accacagagcttaagttcggccataagctcgttaagattgaatggacagatgacaaacaaatttgtcatttcgcaatcggtgaggatctgaaaacgccacacaccgagaaatacgacttcgta
attgggtgcgatggtgcatactcggcgacacgctctcaaatgcagcgtaaagtggaaatggatttctcccaagaatatatgaatcttcggtacatcgagctttacatcccgccgacggaggaat
ttaaaccgaactacggcgggaactttgcaattgctccagatcacttgcacatttggccacggcataaatttatgcttatcgcgctcgccaattccgatgggagctttacgtccacgtttttggtagt
aaagatcagattagtgatttgattacaagtaagtcccgcgtgcgcgagttcttaatcgagaatttcccagacatcattaatatcatggatcttgatgatgctgtaaagcgcttcattacatatccgaa
agaatctctggtatgcgttaattgcaagccatacgatgtcccaggtgggaaagctatcctgctcggtgatgcagcccatgcgatggtgccgttctatgggcagggcatgaattgcggctttgaa
gacgtgcgtattttaatggccctccttaagaaacacagcggtgaccgcagtcgtgcttttactgaatatactcagacgcgccataaagatctcgtatcgattaccgaacttgcgaagcggaacta
caaagaaatgtctcacgacgtgacaagcaagcgcttcttgctgcgtaaaaaacttgatgctttgttctcgattatcatgaaagataaatggatcccgctttataccatgatctccttccgttcggac
attagctactcgcgtgcgctcgagcgcgctgggaagcagacgcgcatcctgaagtttctcgagtccctgaccctgggtatgttatctattggtggttacaagttatttaaatttttgacgcgggag
cgctcatga

**Codon-optimized *hsKMO* from *Homo sapiens***

atggactcttccgtcatccagcggaagaaggtggcagtaattggcgggggtttggtagggagtttacaagcatgctttcttgctaagcgtaattttcaaatcgacgtttacgaggctcgcgaaga
cacccgtgtcgccacttttacacgcgggcgttcaatcaacttagctctgtcccatcgcggtcggcaggcgctgaaagccgtcgggctggaggatcaaatcgtaagtcagggcatccctatgc
gggctcgtatgattcattccttatctgggaagaagtcggcaatcccttatggtacgaaaagtcagtatattttgtcggtctcgcgcgagaatctgaacaaagacctgctcacggcagcggagaa
atatccgaacgtcaaaatgcacttcaaccaccgtttattaaagtgtaacccagaagaaggtatgattactgtccttggctcagacaaagtacctaaagacgttacatgtgatctcatcgttggctgt

gacggtgcatattccacagttcgctcacacctcatgaagaaaccacgcttcgattatagccagcaatacattcctcatgggtacatggagctgacaattcctccaaaaaatggtgattacgccat

ggaaccgaactacttacacatttggcctcggaacacctttatgatgatcgccctccctaacatgaataagtctttcacctgtaccctgtttatgccattcgaagagttcgaaaaattactgacgtcta

atgatgtggtggatttcttccagaaatactttcctgatgctattccgcttattggcgaaaagttgctggttcaggacttcttcttgctgccggctcagccgatgatttctgtcaagtgtagttcctttcat

tttaagtcgcactgcgttttgctgggggatgctgctcatgcaatcgtcccgtttttggccaaggcatgaacgctggctttgaagactgcctcgtatttgatgaattaatggacaaattctccaatga

cctcagtctgtgccttcctgtattctcgcggctgcgtattccggatgatcatgctatttcggacctctcaatgtataactatattgaaatgcgcgcgcacgttaacagttcctggtttattttccaaaaa

aatatggagcggtttctgcacgcgatcatgccttcgacatttatcccgttgtacactatggtgacgttttcacggattcggtatcacgaagcagttcaacgctggcactggcaaaagaaagtcatt

aataaagggctgtttttctgggcagcctcattgccatttcctcaacatacctgttgatccattatatgagtccacgttccttcctctgtcttcggcggccgtggaactggatcgcgcactttcggaat

acaacctgtttcccggcaaaggctgtcgacagcctggaacagatttctaatttgattagtcgttga

**Codon-optimized *sibL* from S*treptosporangium sibiricum***

atgtcgagcgaattatcggctttagtgccagtactttttggccatgccgcctttcaacagcttaatgccggttgccagctgggtttatttgaattgctgcatgagcgcggtcctctctcggctgaag

aagttgccgacgcactccgtcttccacgccgttctgcagacatcctcttgcttggcacgaccgcattgggtctttcaaccgttacagacggtggttatcggaatggcgctccgatcggtgctgc

attccgggacggtctctggccggtattacgggatatcgtgcaataccaagataagatcgcgtatcagcctgcggcggactacgtagagagcctgcgtacgggtcaaaacgctggcattcgg

catttcccggggacgacacgtgacctgtacagccgccttgctgccgtaccgggtttagaggagctgttttatcgtggtatgcacgcatggagccaattatctaacccggtactgcttgcgcagc

cggacttcacccgcgtccaccgcgtgttagatgtaggtggcggcgacgctgtgaacgctgtggctctcgctcgcgcacatccgagtcttcgggttacggtcctcgatcgcccgggtgcttta

gaagtcgctcggaagaccattgcagaagcggggctggaagaacgcgtgcgcacacatgccgccgacattttcacagattcctaccctgcaggtcacgattgcgtcctgttcgcgcatcaac

tggttatttggtcaccagaacagaatctgacgttgctgcgtaaagcatacgatgcagtcgaaccagggggcgtgttctggtgttcaatgccttcactgacgatgaccgtacggggccattata

| |
|---|
| tgccgcattggataatgtttattttacgactttgccttttcgtcacagtaccatccatcggtgggccgactgcgagtcgtggctgcgcgaagcaggctttacagatgtgggtcgcacagcacctc<br>caggttggactcctcacggggttgtgagcggctcgcgtccacggtga |
| **Codon-optimized *sibQ* from *Streptosporangium sibiricum***<br>atggaatctacacgtggctgggcagaaaaggaagcgtctgccgaccctctcgcagttttcgctaaggacttccttatcccagaggggaccttggccttgaacggtaattctctggggccgcct<br>gctgcagctgttccgattgccttgtccaatgcagtacgccaatggtctgagcacttaatccgggggtggtgggacgatgggtggtgggatgcccctgaacgcgtgggtgaccggatcggtc<br>gcctcatgggggctggtccgggccaagtagtagttggtgagtccgcgtctgttcaattgttcaacgctttgacagctgctgtgcgtttagtccctgaacggcgtgtgttaattgcagatgctggg<br>aattttccaaccgatcgttatcttgctgccagcgttgcgcgtcttcttgggctgcgtctggtcgaaacgagcatgaacgacttacctggcgctttggcccgtcatgcagggcaagttggtgcagtt<br>ttatgcggggctgtagattttcgtactggtgagctttgggacgtgccggggctcactacagcgatccatcaagcgggtggggtttcagtatgggatttatcgcacgctgccggtgcagtaccg<br>gtgcaagccgacgccgccggtgttgacctggcagtaggttgcgggtacaagtacttaagtggcggtcctggggcaccagcgtttctgtacgccgcgcggcggcatcaccatgcgcttgatt<br>ttgctgtgacagggtggcacgggcacgcggatccttttgccatgtctgctagtttcgtaccggcggatgggatcacgcgcgcacgcaccgggactcctcagattttgagtcttcttgccctcga<br>agcagcgttagagccgttggaggcagctggtatggcggctgtgcgtgcaaaaagtttgcgcctcgcagattaccttttcgccctcgtggctgatctcgatagcgtcgaaattgtgacgcctcg<br>ggacccgcatcgtcggggtaaccacgtaacattacgcttaccagatgcagtcgcagttgctgaaagtctggcgcgtcggggggttattgttgatgagcgtccaccggacttgttgcgcatttgt<br>ctgaatggcctttacatctcctggactgatgtttatgacgccgtgacacaccttcgggaggttctgcgttga |
| **Codon-optimized *mup1* gene from *Saccharomyces cerevisiae***<br>atgtcagaggggcggaccttcctgtcacagctcaacgtgttcaacaaagagaattatcagttttcttcttccacgaccaaaaaggaggtgtcaaactccaccgtagatgcggacaatggtgctt<br>ctgactttgaagcagggcagcagttcgcaaccgagcttgaccaaggtgaaaagcaactcggcattctcagttgcattggcctcatttgcaatcgcatgttaggtactggcgttttcgcggtttcg |

tcaactatctacacattgtgcgggagcgtaggtttagcattgattatgtgggccgtaggtgcgatcattgcgatcagtggcctttatgtatatatggaattcggtaccgcaattccaaagaatggtg
gcgagaagaattacttggaggcgatttttcgtaagccgaaatttttcatcacgtgcatgtacgcggcatatattttctttcttggttgggctgcgggcaactccatcaacactgcaattatgtttctca
ctgcggcagacaccgaagtgactaaatggaatcaacgggggatcggcgtggcggtagtctttttcgcgttcctcattaatagcctgaacgtaaagatcggtctttacttacagaacatcttgggt
atcttcaagatcgggatcgtgttgttcatctctattactggctgggtagcgctcggtgggggtctgaaggatggctatcaatctcacaactttcggaatgcattcgagggcactgagactgcgac
agcctatggcattgtaaacgctctctattcggttatttggtctttcgttgggtatagcaatgtcaactacgcactgggggaagttaagaatccggtacggactctcaagattgcgggtccgactag
catggtgttccttgcaattatctacatctttgtcaacattgcttatttcgctgtagttccaaaggacaaattgatctcgtcaaaattgatccttgctgctgatttctttgatattgtgtttgggggccaggct
aaacgcgccgctgcggcactggtcgggctttcggcccttggcaatgttctgtcggttatttttctcaagggcgcattattcaacaacttgggcgggaaggtgtattacctttcagtaactttttgc
gtcctcgaaaccattcaattccccgatggtagggttattccagcatttcattgtttgtacagtaaccattcttgcaccgccaccaggcgacgcatatcttttagtgcaaaacttaatttcttaccctatg
aatattatcaattttgcaattagtgccgggttgttgtggatctattggcaacgtcgtcaggggaaaatcgagtggaaccctcctattaaggctggcgtgttcgttactgggtttttacacttagtaat
ctgtacttgattatcgccccatacgtgccaccttccaacggtgaatcagtttactcgtcaatgccgtattggattcattgtgtgatcgcctggggcattttcttctttgggggtgtatactacgttgtct
gggcacaattattgcctcgctggggccactataagctggtttcgaaagatgtactcggcgaggacgggttttggcgtgtgaagatcgctaaggtgtatgatgacacgattggggatgttgatac
tcaagaagacggggtcatcgagacgaatatcattgaacactacaagtctgaacaagagaaatcactgtaa

**Supplementary Table 5 | Relevant studies reporting the microbial production of molecules associated with kynurenine pathway.**

| Product | Producing host | Titer | Cultivation mode | Reference |
|---|---|---|---|---|
| Kynurenic acid | *Yarrowia lipolytica* | 17.7 mg/L | From supernatant after 150 h of cultivation with precursor supplementation using stirred-tank reactor | 4 |
| Quinoline scaffolds | *Saccharomyces cerevisiae* | 1.6 µg/L | Bioreactor conical tubes | 5 |
| Cinnabarinic acid | *Pseudomonas chlororaphis* GP72 | 136.2 mg/L | Fed-batch fermentation using shake flasks | 6 |

#### Supplementary Figures

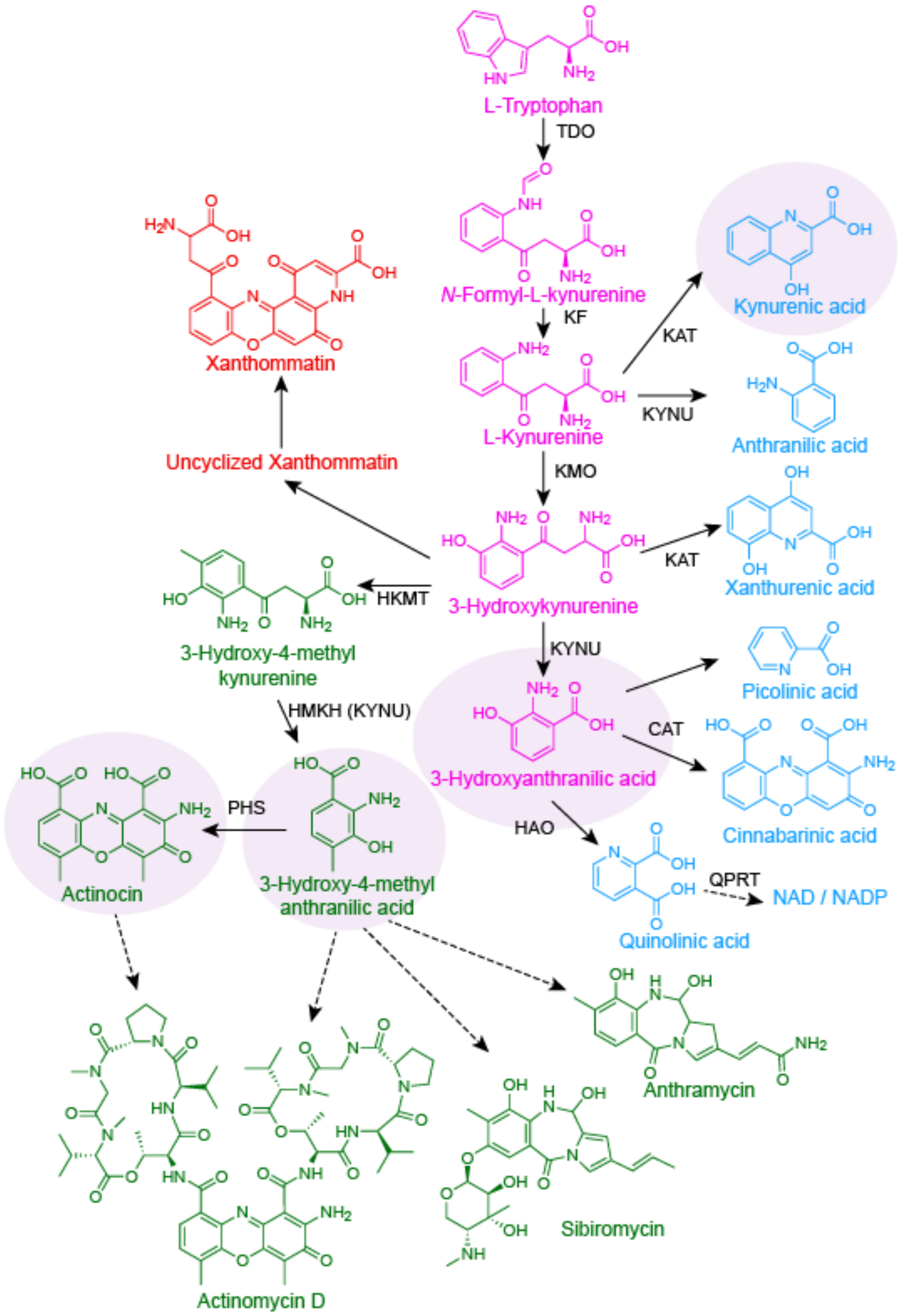


**Supplementary Fig. 1 | Kynurenine pathway leading to the biosynthesis of multiple medicinal molecules in diverse organisms.** Conserved reactions across different organisms are shown in pink, and organism-specific reactions are shown in different colors: red, invertebrates, blue, *Saccharomyces cerevisiae*; and green, *Strepromyces* species.

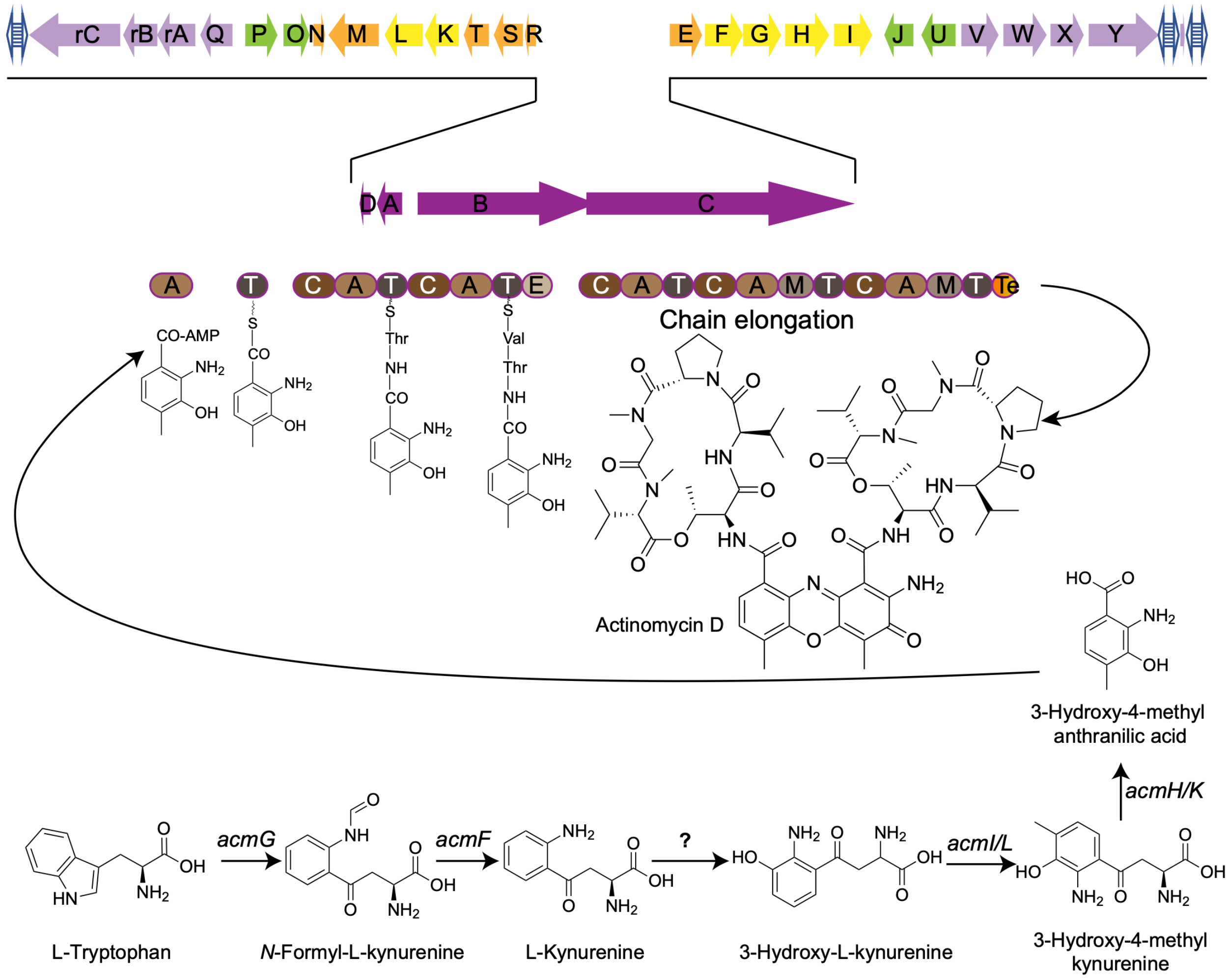


**Supplementary Fig. 2 | Biosynthetic gene cluster of actinomycin D and its corresponding biochemical reactions in *Streptomyces anulatus*.**

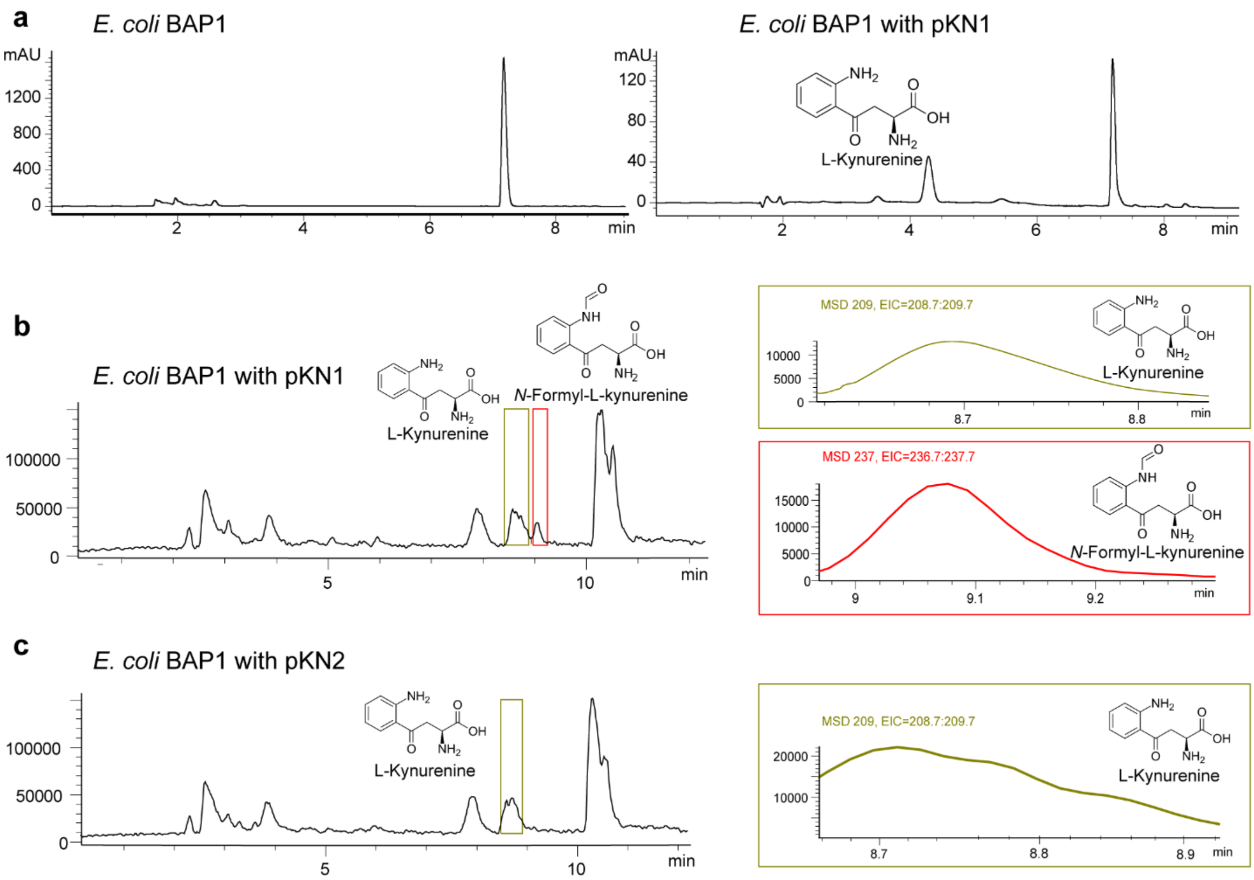


**Supplementary Fig. 3 | HPLC and LC-MS profiles for L-kynurenine and *N*-formyl-L-kynurenine. a,** HPLC profile showing the production of L-kynurenine from *E. coli* BAP1 with pKN1. **b,** LC-MS profile indicating the production of L-kynurenine (green boxes) and its precursor *N*-formyl-L-kynurenine (red boxes) in *E. coli* BAP1 with pKN1. **c,** LC-MS profile indicating the production of only L-kynurenine (green boxes) in *E. coli* BAP1 with pKN2. The precursor *N*-formyl-L-kynurenine was not detected in this case. m/z of L-kynurenine and *N*-formyl-L-kynurenine are: 208.22 g/mol and 236.224 g/mol, respectively.

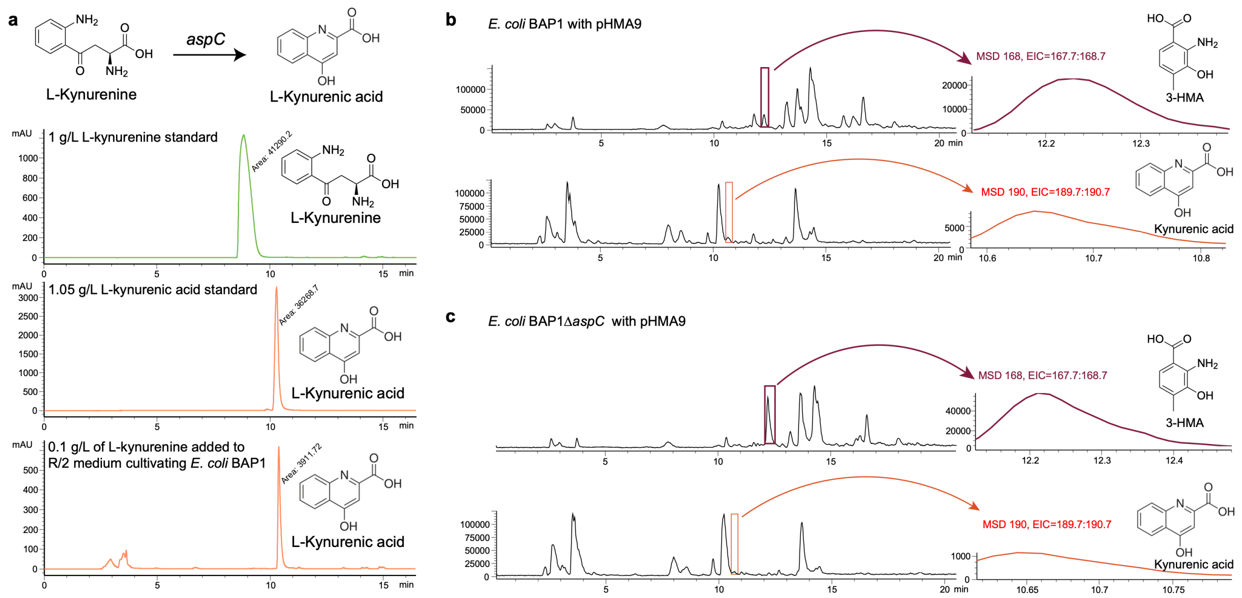


**Supplementary Fig. 4 | HPLC and LC-MS profiles showing the presence of kynurenic acid under three different conditions. a,** HPLC profiles for kynurenic acid by HPLC after 24 h of flask cultivation with 0.1 g/L of L-kynurenine in R/2 medium for *E. coli* BAP1. This shows the additional function of aspartate aminotransferase (encoded by *aspC* gene) converting L-kynurenine to kynurenic acid, as previously reported[7]. **b**, LC-MS profiles showing the production of kynurenic acid along with 3-hydroxy-4-methyl anthranilic acid (3-HMA) in *E. coli* BAP1 with pHMA9. **c,** LC-MS profiles showing the decrease in the kynurenic acid production and the increase in the 3-HMA production in *E. coli* BAP1Δ*aspC* with pHMA9. (**b** and **c**) Upper and lower profiles come from ethyl acetate extracts and supernatants of the same samples, respectively. m/z of 3-HMA and L-kynurenic acid are 167.16 and 189.1675, respectively.

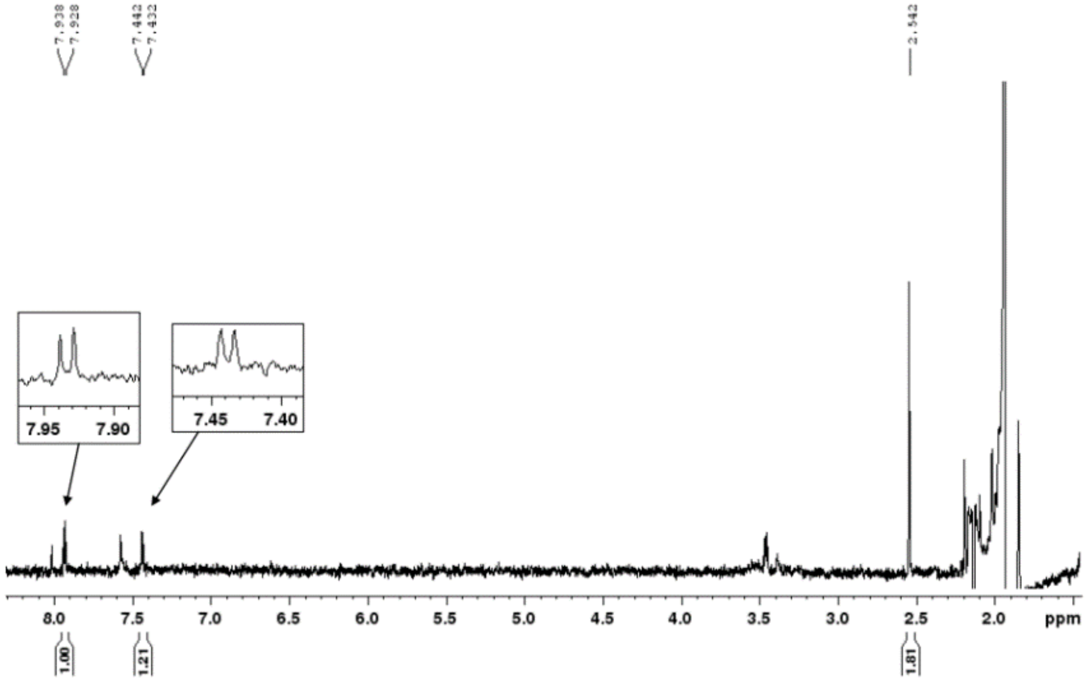


**Supplementary Fig. 5 | Confirmation of actinocin produced from BAP1*ΔaspC* strain with pHMA9, pSAM3 and pACN1 using NMR.**

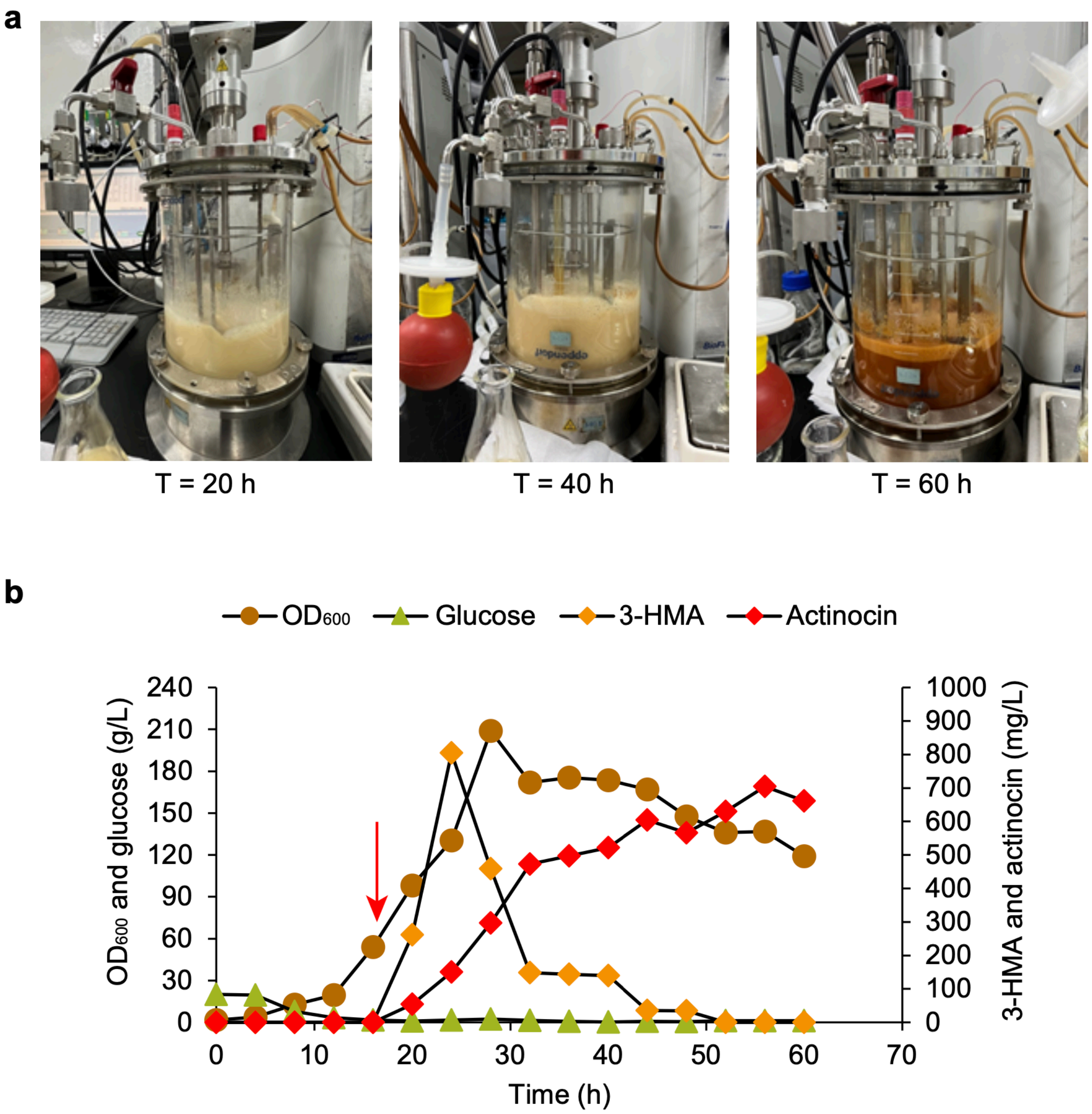


**Supplementary Fig. 6 | Fed-batch fermentation of the final actinocin-producing strain. a,** Fed-batch fermentation progress of the final strain *E. coli* BAP1Δ*aspC*Δ*tnaA* with pHMA9, pSAM4, pTRP1 and pIF at 20 h, 40 h and 60 h. **b,** The second fed-batch fermentation profile of the final actinocin-producing strain confirming the reproducibility of the actinocin production.

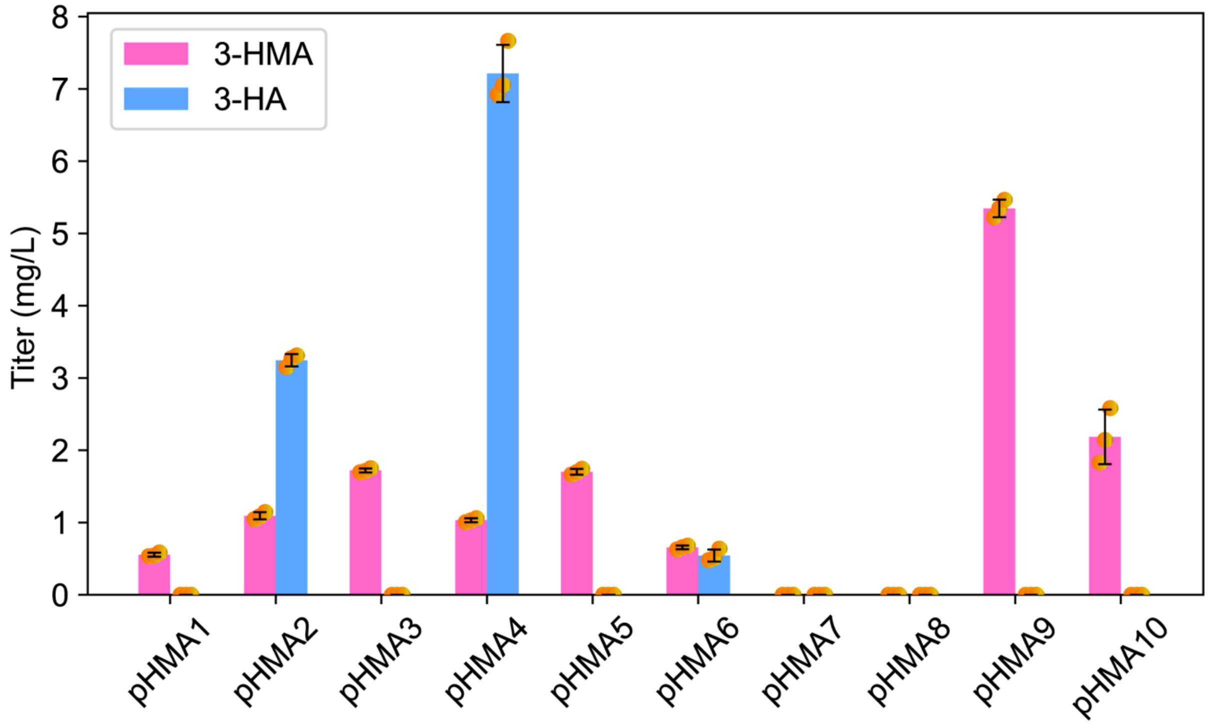


**Supplementary Fig. 7 | Production titers of 3-hydroxyanthranilic acid (3-HA) and 3-hydroxy-4-methylanthranilic acid (3-HMA) from *E. coli* BAP1*ΔaspC* strain harboring different plasmids presented in the graph.** Flask cultivations were performed using R/2 medium supplemented with 5 mM L-tryptophan and 10 mM L-methionine. Data present mean values and their standard deviation from triplicate experiments of flask cultivation.

### Supplementary References


1. Pfeifer, B., Hu, Z., Licari, P. & Khosla, C. Process and metabolic strategies for improved production of *Escherichia coli*-derived 6-deoxyerythronolide B. *Appl. Environ. Microbiol.* **68**, 3287–3292 (2002).
2. Song, C. W. & Lee, S. Y. Rapid one-step inactivation of single or multiple genes in *Escherichia coli*. *Biotechnol. J.* **8**, 776–784 (2013).
3. Du, J., Yang, D., Luo, Z. W. & Lee, S. Y. Metabolic engineering of *Escherichia coli* for the production of indirubin from glucose. *J. Biotechnol.* **267**, 19–28 (2018).
4. Rakicka-Pustułka, M. *et al.* The microbial production of kynurenic acid using *Yarrowia lipolytica* yeast growing on crude glycerol and soybean molasses. *Front. Bioeng. Biotechnol.* **10**, 936137 (2022).
5. Torrens-Spence, M. P., Liu, C.-T. & Weng, J.-K. Engineering new branches of the kynurenine pathway to produce oxo-(2-aminophenyl) and quinoline scaffolds in yeast. *ACS Synth. Biol.* **8**, 2735–2745 (2019).
6. Yue, S. *et al.* Synthesis of cinnabarinic acid by metabolically engineered *Pseudomonas chlororaphis* GP72. *Biotechnol. Bioeng.* **116**, 3072–3083 (2019).
7. Han, Q., Fang, J. & Li, J. Kynurenine aminotransferase and glutamine transaminase K of *Escherichia coli*: identity with aspartate aminotransferase. *Biochem. J.* **360**, 617–623 (2001).
